# Ependymomas are cancers of the pre-neural crest/roof plate lineage

**DOI:** 10.64898/2026.07.13.736818

**Authors:** Polina Balin, Sachin A. Kumar, Riley Galton, Alberto Delaidelli, Wilda Orisme, Parthiv Haldipur, Ncedile Mankahla, Amr Saadeldin, Nik von Krosigk, Maria C. Vladoiu, Anders W. Erickson, Meyer Barembaum, Jake Millman, Amy J. Banks, Jeffrey T. Joseph, Oman Khan, Simon Du, Olga Sirbu, Winnie Ong, Damien Faury, Vicente Santa-Maria Lopez, Sigourney Bonner, Jennifer Coleman, Jason Eigenbrood, Elizabeth A. Cooper, Jonathan K. Duh, Jiao Zhang, John J.Y. Lee, Alexandra Rasnitsyn, Aidan E. Cedillo, Gabrielle Persad, Liam D. Hendrikse, Olivier Saulnier, Randy Van Ommeren, David Przelicki, Namal Abeysundara, Rachel N. Curry, Hannah N. Ahmed, Raul A. Suarez, Cory M. Richman, Ning Huang, Hao Wang, Haipeng Su, Jonelle Pallotta, Esta Mak, Andrew Bondoc, Adrian Levine, Baojin Yao, Laura M. Prolo, Craig Daniels, Andrey Korshunov, Kristen W. Yeom, Vijay Ramaswamy, Livia Garzia, Ana Nikolic, Daniel Schramek, Qing Richard Lu, Jeremy N. Rich, Kenneth Aldape, Kathleen J. Millen, Poul H. Sorensen, Marco Gallo, Richard J. Gilbertson, Nada Jabado, Xiaochong Wu, Lincoln D. Stein, David W. Ellison, Marianne E. Bronner, Michael D. Taylor

**Affiliations:** The Arthur and Sonia Labatt Brain Tumour Research Centre, The Hospital for Sick Children, Toronto, ON, Canada; Developmental and Stem Cell Biology Program, The Hospital for Sick Children, Toronto, ON, Canada; Department of Pediatrics, Baylor College of Medicine, Houston, TX, United States; Texas Children’s Cancer and Hematology Center, Houston, TX, USA; Department of Pediatric Oncology, Dana-Farber Cancer Institute and Division of Hematology/Oncology, Boston Children’s Hospital, Harvard Medical School, Boston, MA, USA; Division of Biology and Biological Engineering, California Institute of Technology, Pasadena, CA, United States; Stowers Institute for Medical Research, Kansas City, MO, United States; Department of Pathology, Mass General Brigham, Harvard Medical School, Boston, MA, USA; Department of Molecular Oncology, British Columbia Cancer Research Centre, Vancouver, BC, Canada; Department of Pathology, St. Jude Children’s Research Hospital, Memphis, TN, United States; Center for Integrative Brain Research, Seattle Children’s Research Institute, Seattle, WA, United States; Development, Disease Models & Therapeutics Graduate Program, Baylor College of Medicine, Houston, TX 77030, USA; Department of Molecular Genetics, University of Toronto, Toronto, ON, Canada; Department of Biochemistry and Molecular Biology, Cumming School of Medicine, University of Calgary, Calgary, AB, Canada; Charbonneau Cancer Institute, Cumming School of Medicine, University of Calgary, Calgary, AB, Canada; Department of Pathology and Laboratory Medicine, Cumming School of Medicine, University of Calgary, Calgary, AB, Canada; Hotchkiss Brain Institute, Cumming School of Medicine, University of Calgary, Calgary, AB, Canada; Alberta Precision Laboratories, Calgary, AB; Department of Medical Biophysics, University of Toronto, Toronto, ON, Canada; The Research Institute of the McGill University Health Centre, Montreal, QC, Canada; Department of Pediatrics, McGill University, Montreal, QC, Canada; CRUK Cambridge Institute, Li Ka Shing Centre, Robinson Way, Cambridge, UK; Stanford University School of Medicine, Palo Alto, CA, United States; Department of Pathology and Krantz Family Center for Cancer Research, Massachusetts General Hospital and Harvard Medical School, Boston, MA, United States; Broad Institute of Harvard and Massachusetts Institute of Technology (MIT), Cambridge, MA, United States; University Health Network, Princess Margaret Cancer Centre, Toronto, ON, Canada; Genomics and Development of Childhood Cancers, Institut Curie, PSL University, Paris, France; INSERM U830, Cancer Heterogeneity Instability and Plasticity, Institut Curie, PSL University, Paris, France; SIREDO: Care, Innovation and Research for Children, Adolescents and Young Adults with Cancer, Institut Curie, Paris, France; Cell Biology Research Program, The Hospital for Sick Children, Toronto, ON, Canada; Department of Pediatric Laboratory Medicine, Hospital for Sick Children, Toronto, ON, Canada; Department of Neurosurgery, Stanford University School of Medicine, Palo Alto, CA, United States; Neuropathology, German Consortium for Translational Cancer Research (DKTK), German Cancer Research Center (DKFZ), Heidelberg, Germany; Department of Radiology, Phoenix Children’s Hospital, 1919 E Thomas Rd, Phoenix, AZ, United States; Division of Haematology/Oncology, Department of Pediatrics, The Hospital for Sick Children, Toronto, ON, Canada; Cancer Research Program, The Research Institute of the McGill University Health Centre, Montreal, QC, Canada; Centre for Molecular and Systems Biology, Lunenfeld-Tanenbaum Research Institute, Mount Sinai Hospital, Toronto, ON, Canada; Brain Tumor Center, Division of Experimental Hematology and Cancer Biology, Cancer and Blood Diseases Institute, Cincinnati Children’s Hospital Medical Center, Cincinnati, OH, USA; Lineberger Comprehensive Cancer Center, University of North Carolina, Chapel Hill, NC, USA; Department of Neurology, University of North Carolina, Chapel Hill, NC, USA; Laboratory of Pathology, National Cancer Institute, Bethesda, Maryland, USA; Department of Pediatrics, University of Washington, Seattle, WA, United States; Department of Pathology and Laboratory Medicine, University of British Columbia,Vancouver, BC, Canada; Alberta Children’s Hospital Research Institute, Cumming School of Medicine, University of Calgary, Calgary, AB, Canada; Dan L Duncan Comprehensive Cancer Center, Baylor College of Medicine, Houston, TX, USA; Adaptive Oncology, Ontario Institute for Cancer Research, Toronto, ON, Canada; Department of Neurosurgery, Baylor College of Medicine, Houston, TX, USA; Department of Neurosurgery, Texas Children’s Hospital, Houston, TX, USA

## Abstract

Distinct molecular variants of the brain cancer ependymoma are distributed along the rostral-caudal extent of the central nervous system (CNS). Historically proposed to arise from ventricular ependyma, recent studies have suggested conflicting cellular origins, including the neural radial glia and the roof plate lineages. Using single-cell transcriptomics, immunohistochemistry, and lineage tracing, we demonstrate that ependymomas across all CNS compartments transcriptionally mirror MSX1^+ve^ pre-neural crest/roof plate (Pre-NC/RP) lineage derivatives. Ependymoma subgroups recapitulate the spatial and molecular diversity of regional Pre-NC/RP populations, while retaining conserved MSX1 expression. Expression of the oncogenic fusion ZFTA-RELA within the murine Pre-NC/RP lineage generated tumors that faithfully resembled human ependymoma. These findings identify a common embryonic cellular origin for ependymomas and reconcile previously conflicting models of tumorigenesis.

## Main

Ependymomas constitute a family of cancers distributed along the rostro-caudal extent of the central nervous system (CNS). Initially thought to represent a single disease due to their similar histology, we now recognize ten clinically and molecularly distinct subgroups arising at discrete anatomic levels of the CNS^1,2^. Most ependymomas are diagnosed in children or young adults, with a minority of cases found in adults. The most common type of ependymoma, **<u>P</u>**osterior **<u>F</u>**ossa **<u>A</u>** (PFA-EPN) is a malignancy of babies and toddlers, with a long-term overall survival of only 50%^1^. Current therapy is maximal safe surgical resection followed by external beam radiation^3^. Neither cytotoxic chemotherapy, nor targeted therapies, are currently standard of care, with limited impact despite numerous clinical trials^4^.

The name ‘ependymoma’ originated in the 1920s, when early histological studies identified ependymal rosettes, formed by ciliated cells surrounding a hollow core^5^. The recognition decades later that ependymal cells are terminally differentiated post-mitotic cells made this an unlikely cell of origin for a cancer^6^. In the current WHO classification, ependymomas are classified as glial tumors, presumed to arise from cells of the neural tube. Indeed, we previously published that ependymomas contain BLBP^+ve^ ‘radial glia-like cells’, that function as cancer stem cells *in vitro^7^*. Subsequent single cell transcriptomics validated the presence of BLBP^+ve^ radial glial-like cells (‘gliogenic progenitors’ (GPs)), but paradoxically also identified ‘roof plate-like stem cells’ (RPSCs)^8^. Roof plate-like stem cells originate from the neural plate border, a region between the neural plate and non-neural ectoderm, that generates the neural crest and roof plate^9^.

We hypothesized that ependymomas originate from a cell type in early embryonic development, capable of giving rise to both ‘gliogenic progenitors’ and ‘roof plate-like stem cells’. Herein, we show that all ependymoma molecular subgroups transcriptionally mirror the MSX1^+ve^ pre-neural crest/roof plate (Pre-NC/RP) lineage of the dorsal neural tube, which indeed gives rise to both radial glia and roof plate-like stem cells. While all ependymomas are MSX1^+ve^, transcriptional variation between ependymoma subgroups mirrors locally distinct Pre-NC/RP lineage differences along the rostral-caudal axis of the CNS. Expression of a ZFTA-RELA transgene under a Pre-NC/RP specific promoter in mice generates ependymomas *in vivo*. Our findings resolve the prior paradox between neural plate and roof plate origins, by demonstrating that ependymomas are not gliomas, but rather cancers of the embryonic dorsal midline.

### PFA-EPN exhibits a stem cell hierarchy

We generated a single cell transcriptomic (scRNAseq) dataset of 41,844 cells from eight primary, therapy-naive PFA-EPN samples, identifying several distinct microenvironmental clusters including microglia, lymphocytes, and vasculature. PFA-EPN neoplastic cells (*SOX2^+ve^, GFAP^+ve^*) were identified based on the presence of somatic copy number variations (**Fig. 1A, fig. S1A-C**). Re-clustering of isolated tumor cells identified seven heterogenous clusters, including neural stem cell-like (NSC), neuronal progenitors (NP), astrocytes (ASTRO), immune reactive cells (IRC), oligodendrocyte precursors (OPC), mesenchymal cells (MC), and ciliated cells (CC) (**Fig. 1B-C, fig. S1D, table S1**). There was no significant difference in the abundance of various cell types across PFA samples (**fig. S1E**). To determine the hierarchical relationship between the various tumoral clusters, we used CytoTRACE, which positioned the NSC cluster at the apex of the tumor hierarchy. Subsequent pseudotime analysis identified three different arms of cellular differentiation (**Fig. 1D**, **fig. S1F**). These results place MSX1^+ve^ ‘neural stem cells’ at the apex of the PFA hierarchy.

**Fig. 1.**
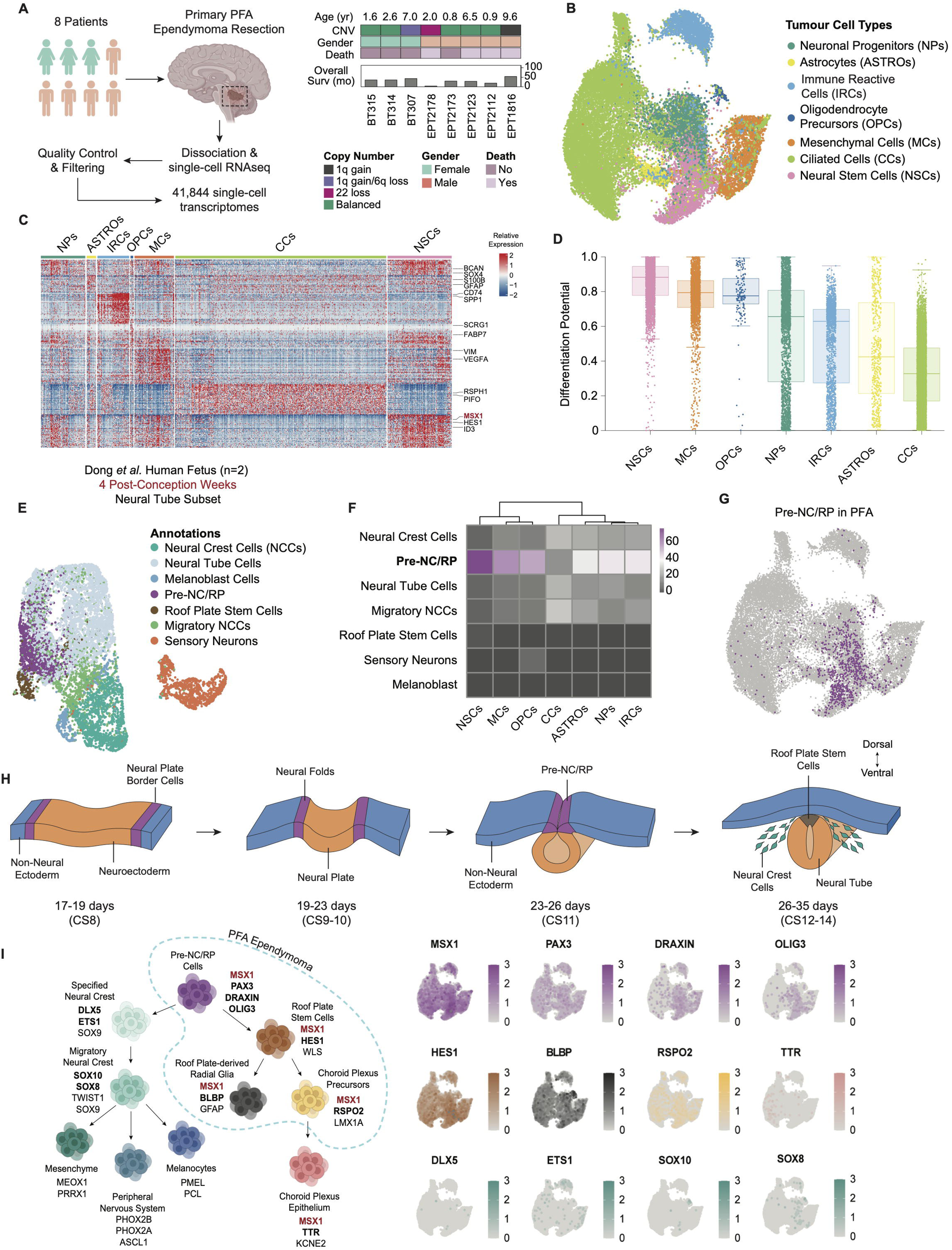
PFA-EPN mirrors the roof-plate arm of the Pre-Neural Crest/Roof Plate (Pre-NC/RP) lineage. A. Diagram of single-cell RNA sequencing (scRNAseq) workflow for primary PFA ependymoma (PFA-EPN) samples (n=8). Heatmap summary demonstrates clinical and molecular characteristics of each sample. B. Uniform manifold approximation and projection (UMAP) embedding of 23,839 PFA-EPN tumor cells identifies seven unique cell types. Clusters were manually annotated based on their transcriptional signatures and previously published markers. C. Heatmap showing relative expression of the top 50 differentially expressed genes defining each tumor cluster. At least two marker genes are highlighted per tumor cell type. Scale bar represents relative expression with set scale -2 to 2. D. Boxplot of tumor clusters ordered by differentiation potential inferred from CytoTRACE. Centre lines show median, box limits indicate interquartile range (IQR), and lower and upper whiskers extend 1.5× the IQR. E. UMAP embedding of neural tube related cell types derived from 4 post-conception weeks (4pcw) human embryos (n=2) reconstituted from Dong *et al*. F. Mapping of individual cells from PFA-EPN scRNAseq demonstrates resemblance to Pre-NC/RP cells. Heatmap shows the fraction of PFA-EPN cells that mapped to each human fetal cluster as inferred by CHETAH, by cell type annotation. Scale bar is percent fraction 0-100%. G. UMAP embedding of PFA-EPN tumor cells, highlighting cells enriched for a Pre-NC/RP signature. H. Illustration of neural tube closure, outlining differentiation of neural plate borders from Carnegie stage 8 – 14. I. The Pre-NC/RP normal developmental hierarchy (left) with published marker genes. UMAP embeddings (right) of PFA-EPN scRNAseq demonstrating expression of select marker genes for Pre-NC/RP cells (purple), roof plate stem cells (brown), roof plate-derived radial glia (black), choroid plexus precursors (yellow), choroid plexus epithelium (green), and pan-neural crest (turquoise). Normalized set-scale 0-3.

### PFA-EPN mirrors the Pre-NC/RP roof plate arm

To identify an earlier embryonic cell type that serves as a precursor to both GPs and RPSCs, we generated a scRNAseq dataset from four E9.5 murine hindbrains, identifying 19 unique clusters, including RPSCs (**fig. S2A-B, table S2**). Comparison of human PFA-EPN to the E9.5 murine hindbrain demonstrated strong transcriptional similarity between the apical PFA-EPN cluster (NSCs) and murine Pre-NC/RP cells (*Msx1^+ve^, Pax3^+ve^, Olig3^+ve^*) (**fig. S2C**). Due to known stark cross-species differences between the embryonic human and murine hindbrain, and their relevance to hindbrain neoplasia, we re-analyzed published scRNAseq data from two 4 post-conception weeks (4 PCW) human embryos (approximately Carnegie Stage 12 (CS12))^10–12^. Both Pre-NC/RP (*MSX1^+ve^, DRAXIN^+ve^, OLIG3^+ve^*), and RPSCs were present in this human fetal dataset (**Fig. 1E, fig. S3A-B**). Consistent with the mouse hindbrain, comparison to the human fetal neural tube demonstrates high transcriptional similarity between the apical PFA-EPN cluster (NSCs) and the CS12 Pre-NC/RP cluster (**Fig. 1F**). A transcriptional signature derived from the top 70 differentially expressed genes in CS12 human embryonic Pre-NC/RPs was subsequently used to annotate these stem cells in our PFA-EPN single-cell dataset (**Fig. 1G, table S3**). To validate these findings, we used previously published ependymoma single cell data and again found the presence of a Pre-NC/RP transcriptional signature (**fig. S3C-D**)^13^. These findings implicate Pre-NC/RPs of the early embryonic neural tube (4 PCW) as a putative cell of origin for PFA-EPN.

In early human development, neural plate border cells are interposed between the medial neural plate (future CNS), and the lateral non-neural ectoderm (future cutaneous epidermis), (**Fig. 1H**)^14^. Growth of the neural plate initiates the folding of the neural tube, with the neural plate borders becoming the neural folds^15^. The neural folds eventually fuse to seal the neural tube and later form the roof plate along the dorsal midline of the developing CNS^16^. Pre-NC/RP cells are first identified within the neural folds^17^. SOX10^+ve^ neural crest cells (NCCs) begin to delaminate from the closing neural tube, migrating ventrally to form a myriad of neural crest derivatives^18^. The remaining cells along the dorsal midline of the neural tube subsequently differentiate into RPSCs, which eventually give rise to the choroid plexus^16,18,19^.

PFA-EPN tumors express transcriptional markers of Pre-NC/RPs (*MSX1^+ve^*, *DRAXIN^+ve^*, *PAX3^+ve^*, *OLIG3^+ve^),* roof plate-like stem cells (*MSX1^+ve^, HES1^+ve^, WLS^+ve^*), roof plate-derived radial glia (RPRG) (*MSX1^+ve^, BLBP^+ve^, GFAP^+ve^),* and choroid plexus precursor cells (*MSX1^+ve^, RSPO2^+ve^, LMX1A^+ve^*) (**Fig. 1I**). *MSX1* is expressed across all PFA-EPN tumor clusters (**Fig. 1I**). However, we did not observe expression of pan-neural crest marker genes (*SOX10*, *DLX5*, *ETS1* or *SOX8),* suggesting that PFA-EPN does not resemble the neural crest arm of the Pre-NC/RP lineage. We also did not observe robust expression of mature choroid plexus markers such as *TTR* or *KCNE2* (**Fig. 1I**).

Next, we employed scRNAseq of lineage tracing using *Msx1^CreERT2^; R26R-lsl-tdTomato* mice, induced at E9.5. We collected the developing hindbrain at E10.5, E14.5, and E18.5, with robust recovery of the various expected cell types **(fig. S4A-C, table S4)**. Traced tdTomato cells arising from the Pre-NC/RP lineage were subset, and PFA-EPN showed strong mapping of tumor cells to Pre-NC/RP cells themselves, with only partial mapping to RPSCs and choroid plexus **(fig. S4D-G)**. These data support a model in which PFA-EPN mirrors the roof plate arm of the Pre-NC/RP lineage, resolving previous paradoxical observations by unifying RPSCs and a subset of GPs as arising from a single Pre-NC/RP progenitor (**fig. S5A, table S5**)^8^.

### MSX1 is a novel biomarker of for ependymomas

MSX1 is a highly conserved transcription factor in chordates, known to be expressed in the dorsal midline of the neural tube, and is retained across all progenies of the roof plate arm of the Pre-NC/RP lineage^20^. Expression of MSX1, as assessed in sagittal sections of the developing *Mus musculus* and *Homo sapiens* hindbrain, was abundant in the cranial mesenchyme, roof plate, and choroid plexus (**Fig. 2A**). Early in development (CS14 in humans, E9.5 in the mouse), MSX1 is also transiently expressed in the upper rhombic lip (URL) (**Fig. 2A**)^21^. We found expression of MSX1 in all 108 PFA-EPN patient samples, with ubiquitous protein expression across all PFA-EPN cells (**Fig. 2B**). Conversely, MSX1 expression was not observed in medulloblastoma or diffuse midline gliomas including the pons (DIPG) (**Fig. 2B**). The choroid plexus is derived from the MSX1^+ve^ roof plate, and as expected choroid plexus tumors express MSX1 (**fig. S6A**). Excitingly, MSX1 is also expressed in every sample of a large panel of ependymoma variants across the supratentorial, posterior fossa (hindbrain), and spinal compartments (**fig. S6B**). Additional staining of PFA-EPN tumors with the Pre-NC/RP marker PAX3 shows unique expression compared to glioma controls (**fig. S6C**). Thus, MSX1 is an excellent biomarker to identify all the members of the ‘ependymoma’ family by immunohistochemistry in a clinical setting.

**Fig. 2.**
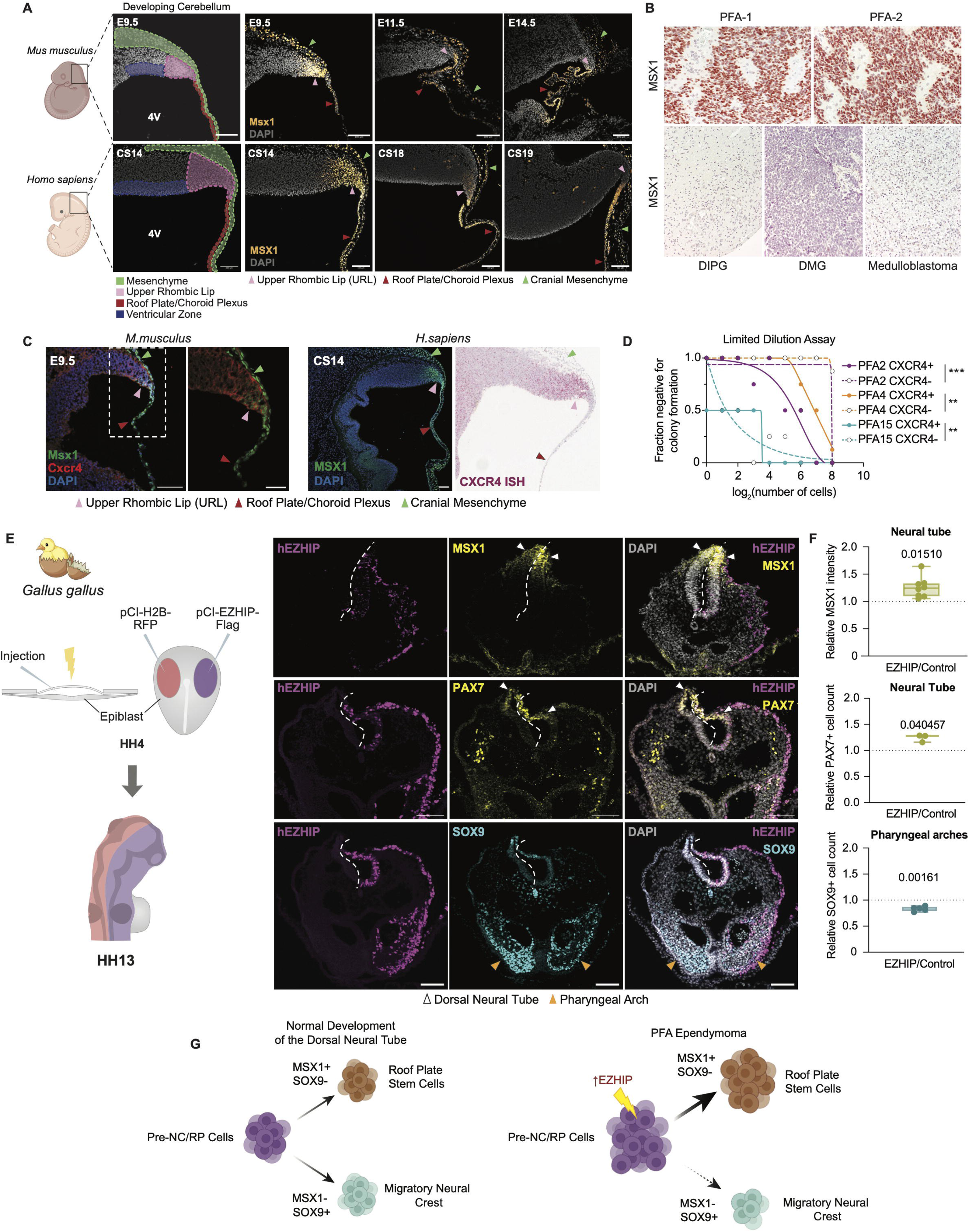
Pre-NC/RP cells recapitulate PFA-EPN biology *in vitro* and *in vivo*. A. Immunofluorescence through sagittal sections of the developing mouse (top) and human (bottom) embryonic hindbrain shows conserved MSX1 expression in the upper rhombic lip, roof plate, choroid plexus, and cranial mesenchyme. MSX1 (yellow) and DAPI (grey). Arrowheads indicate anatomic region. Scale bar is 100μm. 4V = fourth ventricle, CS = Carnegie stage. B. Immunohistochemical staining of MSX1 in patient samples is positive (red) in PFA-EPN tumors, but not negative control diffuse intrinsic pontine glioma (DIPG) and medulloblastoma (MB) tumors. Representative images from both subtypes of PFA-EPN (PFA1, PFA2) are displayed. C. Co-expression of Pre-NC/RPs markers within the upper rhombic limb (URL) using immunofluorescence or *in situ* hybridization in the mouse (left, E9.5) or human (right, CS14) hindbrain. Msx1 (green), Cxcr4 (red), and DAPI (blue). Scale bar 100μm. Arrowheads indicate anatomic region. D. Limiting dilution assay (LDA) demonstrates enrichment of CXCR4^+ve^ colony forming stem cells across three PFA-EPN primary tumor derived cell lines. Statistical comparison performed using Chi-squared test (extreme LDA software), with ** p < 0.01 and *** p < 0.001. E. Electroporation of the *Gallus gallus* embryo with control (pCI-H2B-RFP) or human *hEZHIP-Flag* plasmids prior to neural-tube folding at Hamburger-Hamilton stage 4 (HH4). Representative axial images at HH13, stained for DAPI (grey), hEZHIP (purple), MSX1 (yellow, top), PAX7 (yellow, middle) or Sox9 (cyan, bottom). F. Boxplots showing quantification of relative intensity of MSX1 (top), and relative cell count of PAX7^+ve^ cells (middle) within the neural tube and SOX9^+ve^ cells within the pharyngeal arches (bottom). Arrowheads indicate anatomic structure. Statistical comparison using a two-tailed paired T-test. Centre line shows median and lower and upper whiskers extend 1.5× the IQR. G. Proposed model whereby high expression of EZHIP mediates PFA-EPN oncogenesis in Pre-Pre-NC/RP stem cells by restricting neural-crest migration and persistent accumulation of Pre-NC/RP and roof-plate stem cells.

### Ependymoma derived Pre-NC/RP cells exhibit cancer stem cell properties

PFA-EPN exhibits a steep stem-cell hierarchy, with only a minority of cells capable of tumor repopulation as demonstrated by limiting dilution analysis^22^. Exposure of PFA-EPN cultures to 21% ambient oxygen greatly diminishes their growth potential, and only growth in hypoxia (1% oxygen) permits sustained growth *in vitro^22^*. *CXCR4* is a surface marker expressed in the dorsal midline of the developing human and murine hindbrain, overlapping with MSX1 in the Pre-NC/RP population (**Fig. 2C**)^23^. Limiting dilution analysis (LDA) of CXCR4^+ve^ versus CXCR4^-ve^ flow-sorted populations from patient derived PFA cultures demonstrates significantly increased colony forming ability of the CXCR4^+ve^ compartment (**Fig. 2D**). Consistently, exposure of patient derived PFA cultures to normoxia (21% oxygen), but not hypoxia (1% oxygen) depletes the relative abundance of CXCR4^+ve^ cells across a panel of patient derived PFA-EPN cultures (**fig. S7A**). We conclude that Pre-NC/RP cells, marked by CXCR4^+ve^ expression, are found in human PFA-EPN cultures, and exhibit cancer stem cell properties *in vitro*.

### EZHIP stalls differentiation of Pre-NC/RP cells *in vivo*

While most PFA-EPN have no recurrent somatic mutations, a small fraction of tumors harbor mutations in the Polycomb Repressive Complex 2 (PRC2) inhibitor *EZHIP*, or the canonical H3K27M mutation of histone 3.3 found in diffuse midline gliomas^24–26^. Conversely, nearly all PFA-EPN highly express EZHIP. Indeed, loss of staining for H3K27me3 is a clinical biomarker of PFA-EPN^27^. Interestingly, EZHIP was not expressed in the developing human hindbrain in normal development (**fig. S8A)**. Nonetheless, we hypothesized that Pre-NC/RPs would be sensitive to over-expression of EZHIP *in vivo*. To test this, we electroporated human *EZHIP* (pCl-hEZHIP-FLAG) or control (pCl-H2B-RFP) plasmids into opposite halves of the neural tube of *Gallus gallus* embryos at Hamburger-Hamilton Stage 4 (HH4). Embryos were harvested at Hamburger-Hamilton Stage 13 (HH13), and EZHIP expression was confirmed, including functional repression of H3K27me3 (**Fig. 2E, fig. S9A-B**). EZHIP over-expression was associated with increased expression of Pre-NC/RP markers PAX7 and MSX1, and diminished numbers of SOX9 differentiated neural crest cells in the pharyngeal arches (**Fig. 2E-F**). Notably, H3K27me3 marks accumulate in both the mouse and human Pre-NC/RP hierarchy as cells differentiate into the roof plate and then the choroid plexus (**fig. S9C**). These *in vivo* data are consistent with a model whereby inappropriate EZHIP expression in hindbrain Pre-NC/RP inhibits differentiation and thereby impairs delamination of NCCs. Failure of differentiation leads to the accumulation of Pre-NC/RPs, RPSCs, and their derivatives, giving rise to PFA-EPN tumors **(Fig. 2G).**

### PFB-EPN arises from Pre-NC/RP derived midline glia

Pre-NC/RPs are found across all levels of the developing neural tube, and the Pre-NC/RPs marker MSX1 is expressed across all ependymomas. Therefore, we hypothesized that other molecular subgroups of ependymoma also arise from Pre-NC/RPs across the neuraxis. PFA-EPN is histologically identical to **P**osterior **F**ossa **B** (PFB-EPN), and these two ependymoma subgroups can only be mutually distinguished using molecular biology^28,29^. Single nucleus RNA sequencing (snRNAseq) of three primary PFB-EPNs demonstrates a cluster of cells exhibiting a Pre-NC/RP signature (**Fig. 3A, fig. S10A-D, table S6**). Again, all cases of PFB-EPN tumors (n=12) express MSX1 (**Fig. 3B**).

**Fig. 3.**
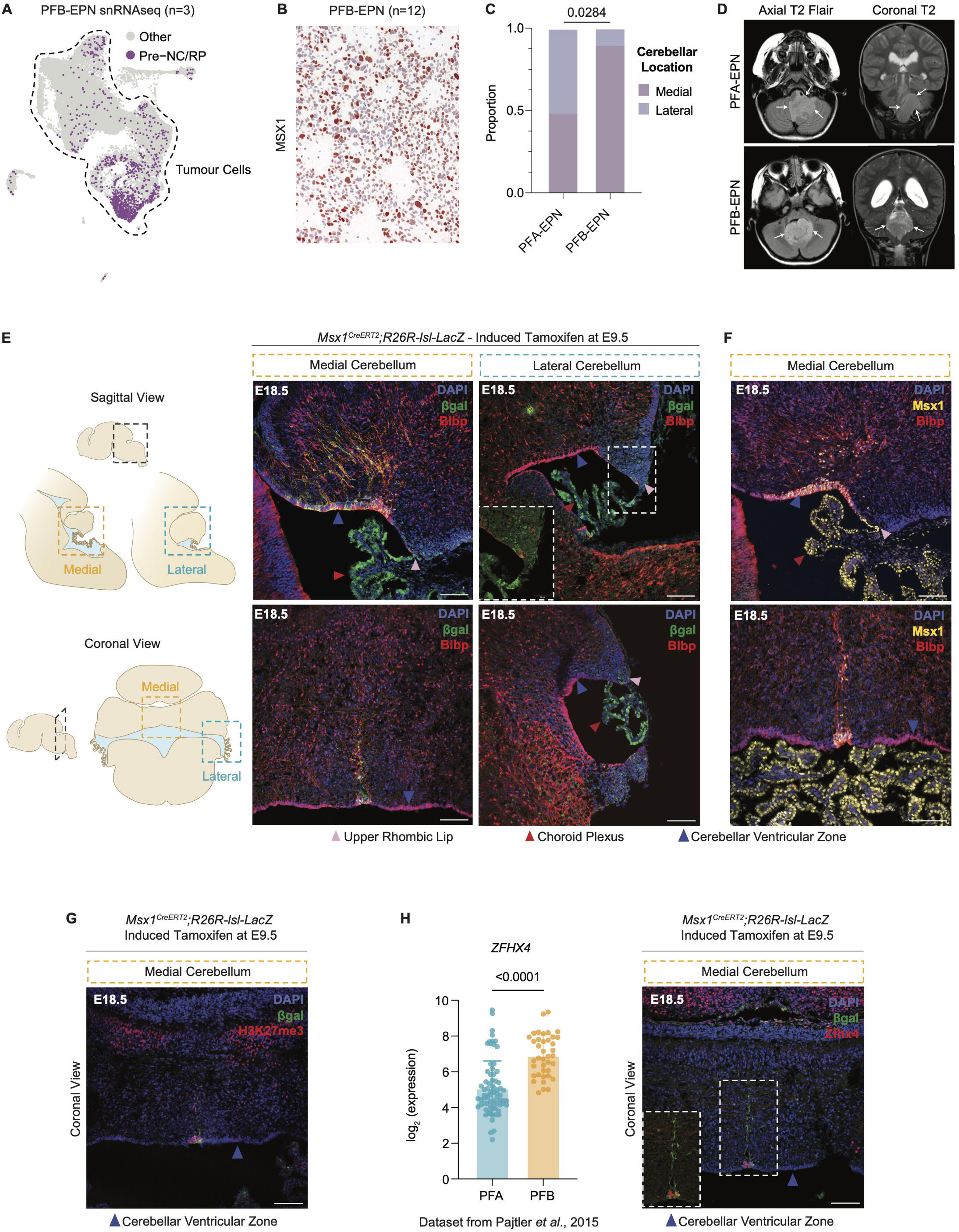
PFB-EPN resembles Pre-NC/RP derived midline radial glia. A. UMAP embedding of PFB-EPN tumor cells (snRNAseq), highlighting cells enriched for a Pre-NC/RP signature. B. Immunohistochemical staining of MSX1 in patient samples is positive (red) in all PFB-EPN tumors (n=12). C. Stacked bar plot comparing the medial versus lateral incidence of PFA- and PFB-EPN tumors within the hindbrain. Statistical comparison performed using Fischer’s exact test. D. Representative axial T2 FLAIR (left) and coronal T2-weighted (right) MRI images of PFA-EPN (top) and PFB-EPN (bottom) tumors. The PFA-EPN tumor is asymmetric with the mass extending into the left perimedullary cistern, surrounding the brainstem and the cerebellum, and inferiorly into the upper cervical spinal canal. The PFB-EPN tumor is well circumscribed and centrally located within the 4^th^ ventricle. E. Lineage tracing of *Msx1^CreERT2^;R26R-lsl-LacZ* mice induced with tamoxifen at E9.5, demonstrates a unique population of Pre-NC/RP derived midline glia, marked by βgal and Blbp co-expression. Sagittal (top) and coronal (bottom) sections through medial (orange) and lateral (blue) hindbrain/cerebella at E18.5 stained for βgal (green), Blbp (red) and DAPI (blue). Scale bar 100μm. Arrowheads indicate anatomic region. F. Pre-NC/RP derived midline glia maintain MSX1 expression at E18.5. Sagittal (top) and coronal (bottom) sections of the medial cerebellum at E18.5, stained for Msx1 (yellow), Blbp (red) and DAPI (blue). Scale bar 100μm. Arrowheads indicate anatomic region. G. Pre-NC/RP derived midline glia express histone-3 lysine 27 trimethylation (H3K27me3). Coronal section of cerebellum at E18.5 in lineage traced *Msx1^CreERT2^;R26R-lsl-LacZ* mice, stained for βgal (green), H3K27me3 (red) and DAPI (blue). Scale bar 100μm. H. The PFB-EPN enriched marker ZFHX4 is uniquely expressed in Pre-NC/RP derived midline glia. Bar plot (left) of ZFHX4 expression in PFB-EPN compared to PFA-EPN from Pajtler *et al.* (2015). Statistical comparison performed using two-tailed unpaired T-test. Coronal section of the cerebellum (right) at E18.5 in lineage traced *Msx1^CreERT2^;R26R-lsl-LacZ* mice, stained for βgal (green), Zfhx4 (red) and DAPI (blue). Scale bar 100μm.

While PFA-EPN is frequently located laterally in the posterior fossa, occupying the Foramen of Luschka, PFB-EPN is almost always midline in the fourth ventricle (**Fig. 3C-D**)^30^. We hypothesized that there exists variability between midline and lateral Pre-NC/RP lineage derivatives within the hindbrain, which produces the clinical, anatomic, and transcriptional differences between PFA-EPN and PFB-EPN. To assess this, we performed lineage tracing in *Msx1^CreERT2^; R26R-lsl-LacZ* mice, with tamoxifen induction at E9.5, corresponding to a developmental stage when Pre-NC/RP cells exist. We harvested embryos at E18.5 and examined the medial and lateral cerebellum in both sagittal and coronal planes. We observe a narrow bundle of cells in the midline of the rostral fourth ventricle which co-expresses both βgal (Msx1) and the radial glial marker Blbp at E18.5, consistent with locally specific radial glia derived from the Pre-NC/RP compartment (**Fig. 3E**). We observe only rare double positive cells in the lateral fourth ventricle at E18.5. These βgal^+ve^/Blbp^+ve^ cells retain high levels of Msx1 expression (**Fig. 3F**).

Clinically, staining for H3K27me3 is used to distinguish PFA-EPN (absent) from PFB-EPN (highly expressed)^28^. We stained medial versus lateral sections for H3K27me3, to determine if this is a retained phenotype. We observed high levels of H3K27me3 within the βgal^+ve^ rostral midline glia, but not in the lateral hindbrain (**Fig. 3G, fig. S10E**). The gene *ZFHX4* is highly expressed in PFB-EPN, distinguishing it from PFA-EPN^1^. We assessed expression of Zfhx4 in the medial versus lateral cerebellum and found that only midline glia were positive (**Fig. 3H, fig. S10F**). This MSX1*^+ve^*/BLBP*^+ve^*population in the dorsal midline of the fourth ventricle occupies the same anatomic location as PFB-EPN, retains H3K27me3, and expresses the marker ZFHX4, making these Pre-NC/RP derived midline glia an excellent candidate for the cell-of-origin of PFB-EPN.

### Spinal ependymomas resemble roof plate-derived radial glia

Ependymomas of the spinal cord comprise multiple distinct entities, most commonly myxopapillary ependymomas (MPE-EPN) of the lower spinal cord (conus medullaris and cauda equina), and intramedullary ependymomas (SP-EPN) of the cervical and thoracic spinal cord^1^. We generated snRNAseq data from three MPE-EPNs and were again able to identify the transcriptional signature of Pre-NC/RPs in tumor cells (**fig. S11A, fig. S12, table S7**). Immunohistochemistry from both MPE-EPN (5/5 cases) and SP-EPN (8/8 cases) tumors demonstrate widespread expression of MSX1 across all tumor cells (**fig. S11B**). *HOXB13*, a marker gene of the caudal spinal cord, is expressed in the tumor cells of MPE-EPN (**fig. S11C**)^31^. Within the dorsal midline of the spinal cord, roof plate cells arising from Pre-NC/RPs transform into a stream of radial glia-like cells, extending processes from the ependyma of the central canal to the pial surface (**fig. S11D**). Lineage tracing in *Msx1^CreERT2^; R26R-lsl-LacZ* mice induced at E9.5 confirms previously published reports of the Msx1 lineage giving rise to Blbp/βgal double positive roof plate-derived radial glia (RP-RG) at E18.5 (**fig. S11E**)^32^. RP-RG of the spinal cord maintain expression of Msx1 and Hoxb13 in the lumbar spine at E18.5, but do not exhibit immunopositivity for oligodendrocyte (Sox10) or neuronal (NeuN) lineage markers (**fig. S11E**). The presence of cells that transcriptionally mirror MSX1^+ve^ roof plate-derived radial glia in both SP-EPN and MPE-EPN is consistent with a model whereby these tumors originate from regional derivatives of the Pre-NC/RP lineage.

### Hindbrain subependymomas transcriptionally mirror the area postrema

Subependymomas are typically indolent tumors, often diagnosed incidentally on neuroimaging or found at the time of autopsy^33^. Like other subgroups, subependymomas also express MSX1 (7/7 cases) (**fig. S13A**). Subependymomas of the fourth ventricle (PF-SE) arise in the subependymal space at the caudal end of the medulla oblongata^34^. This region contains a circumventricular organ called the ‘area postrema’, otherwise known as the ‘chemoreceptor trigger zone’, the vomiting center of the brain^35,36^. Given PF-SEs arise in the area postrema, we hypothesized that Pre-NC/RP cells might be implicated in its development. Indeed, we observe a strong βgal signal (MSX1 lineage) within the area postrema in *Msx1^CreERT2^; R26R-lsl-LacZ* mice induced with tamoxifen at E9.5. Furthermore, a subset of the cells retain Msx1 expression at E18.5 (**fig. S13B**). MSX1 is also expressed in the area postrema of the adult *Homo sapiens* brain (**fig. S13C**). Previously reported marker genes for PF-SE, *KIT* and *GRIN3A*, are also expressed in the *Mus musculus* area postrema (**fig. S13D-E**)^1^. For the first time, we demonstrate that the area postrema is derived from Pre-NC/RP cells, and represents the origin of 4^th^ ventricular subependymomas.

### ZFTA-EPN mirrors the cortical hem lineage

Most supratentorial ependymomas harbour somatic ZFTA-RELA fusions, or other ZFTA fusion oncogenes^37^. scRNAseq analysis of four ependymomas harboring ZFTA fusions (ZFTA-EPN) demonstrates a population expressing the Pre-NC/RP signature (**Fig. 4A, fig. S14A-D, table S8**). All cases of ZFTA-EPNs (16/16) were immunopositive for MSX1 (**Fig. 4B**). In the murine telencephalon, the dorsal midline roof plate invaginates ventrally around E10.5, dividing the neural tube and creating the left and right lateral ventricles. The invaginated roof plate gives rise to the choroid plexus, while the proximal portion, called the cortical hem, gives rise to Cajal-Retzius neurons around E12.5^38,39^. Cajal-Retzius cells migrate to the subpial surface of the cerebral hemispheres, where they provide guidance cues for the laminar organization of cortical neurons ^40^. Later (E13.5-14.5), the cortical hem begins producing fimbrial glia, a unique population of radial glia-like cells adjacent to the nascent hippocampus, before the cortical hem thins (E18.5) and becomes the fimbrial neuroepithelium (**Fig. 4C**)^41,42^.

**Fig. 4.**
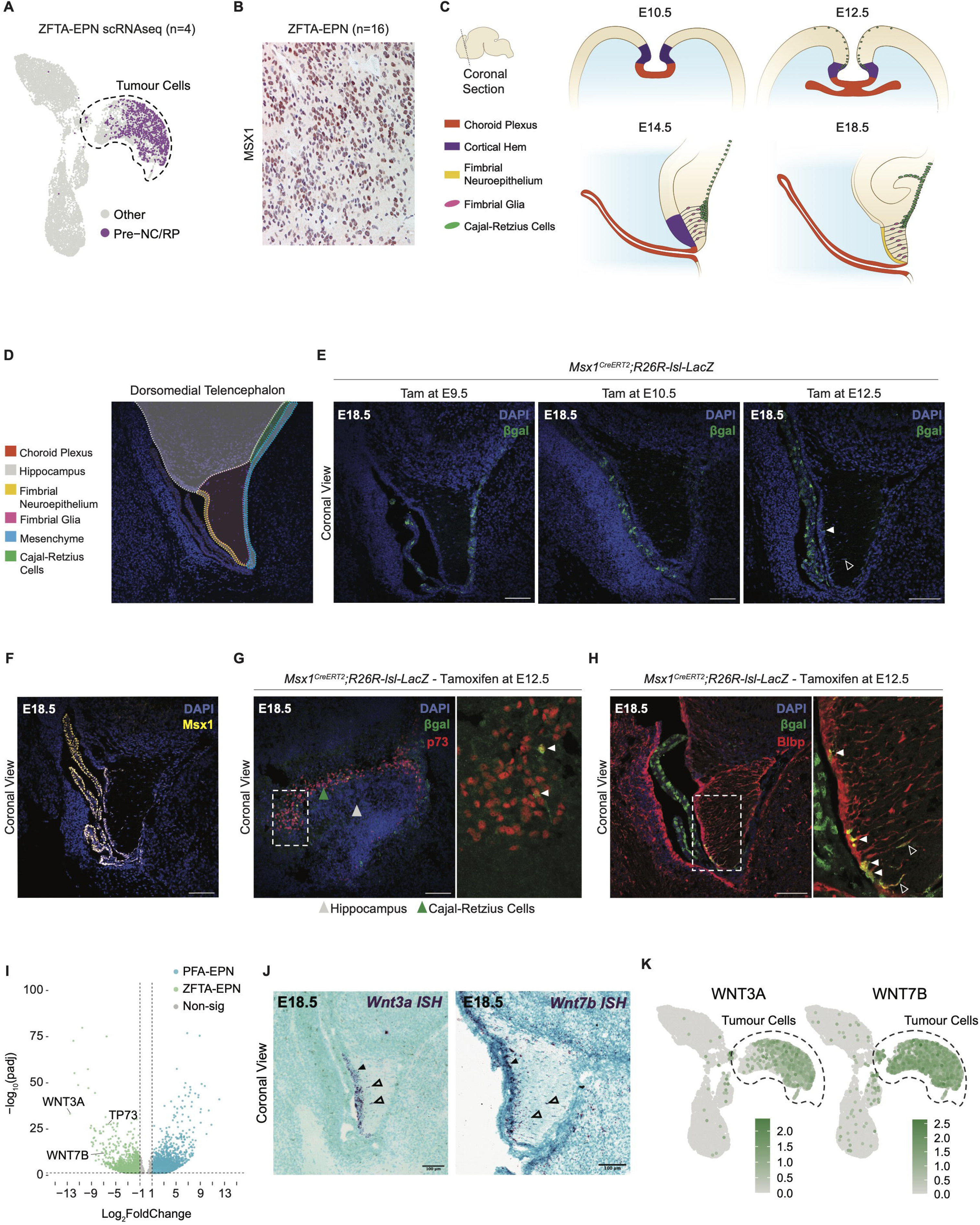
ZFTA ependymoma mirrors the telencephalic cortical hem lineage. A. UMAP embedding of ZFTA-EPN tumor cells (scRNAseq), highlighting cells enriched for a Pre-NC/RP signature. B. Immunohistochemical staining of MSX1 is positive (red) in all ZFTA-EPN tumors (n=16). C. Illustration of coronal sections through the developing murine telencephalon from E10.5 to E18.5, highlighting development of the telencephalic midline. D. Overlay of structures in a coronal section of the E18.5 murine cortical hem stained with DAPI, including choroid plexus (orange), fimbrial glia (pink), hippocampus (grey), mesenchyme (blue), fimbrial neuroepithelium (yellow) and Cajal-Retzius cells (green). Scale bar 100μm. E. The Pre-NC/RP lineage contributes to the developing cortical hem, including the choroid plexus, fimbrial neuroepithelium, and fimbrial glia cells. Coronal sections through the cortical hem at E18.5 in lineage traced *Msx1^CreERT2^;R26R-lsl-LacZ* mice induced with tamoxifen at E9.5 (left), E10.5 (middle), and E12.5 (right). Sections are stained for βgal (green) and DAPI (blue). Arrowheads denote βgal signal within the fimbrial neuroepithelium (filled) and individual fimbrial glia (empty). Scale bar 100μm. F. Cortical hem structures maintain MSX1 expression at E18.5. Coronal section through the cortical hem at E18.5, stained for Msx1 (yellow) and DAPI (blue). Scale bar 100lm. G. Pre-NC/RPs give rise to Cajal-Retzius cells within the hippocampal fissure. Coronal sections through the hippocampus at E18.5 in lineage traced *Msx1^CreERT2^;R26R-lsl-LacZ* mice induced with tamoxifen at E12.5, stained for βgal (green), p73 (red), and DAPI (blue). Arrowheads indicate anatomic region. Inset with arrowheads identify βgal /p73 double-positive Cajal-Retzius cells. Scale bar 100μm. H. Pre-NC/RPs give rise to fimbrial glia cells and contribute to the fimbrial neuroepithelium. Coronal sections through the cortical hem at E18.5 in lineage traced *Msx1^CreERT2^;R26R-lsl-LacZ* mice induced with tamoxifen at E12.5, stained for βgal (green), Blbp (red), and DAPI (blue). Inset with arrowheads identifies βgal/Blbp double-positive fimbrial glia (empty) and fimbrial neuroepithelium (filled). Scale bar 100μm. I. Cortical hem and Cajal-Retzius genes are enriched in ZFTA-EPN compared to PFA-EPN. Volcano plot highlighting cortical hem (*WNT3A, WNT7B)* and Cajal-Retzius (*TP73)* genes. Differentially expressed genes (log_2_FoldChange > 1, P_adj_ < 0.05) indicated in color. J. RNAscope *in situ* hybridization in a coronal section through the murine cortical hem at E18.5, shows *Wnt3a* (left) and *Wnt7b* (right) expression within the fimbrial neuroepithelium (filled arrowheads) and glia (empty arrowheads). Scale bar 100μm. K. UMAP embedding of ZFTA-EPN demonstrates expression of *WNT3A* and *WNT7B* within tumor cells.

We performed lineage tracing in *Msx1^CreERT2^; R26R-lsl-LacZ* mice induced with tamoxifen at either E9.5, E10.5, or E12.5 and harvested the embryos at E18.5. Induction at E9.5 only labelled a few cells within the lateral ventricle (LV) choroid plexus or the cranial mesenchyme. At E10.5, we observed increased recombination within the LV choroid plexus, but no βgal signal within either the fimbrial glia or fimbrial neuroepithelium (**Fig. 4D-E**). In contrast, at E12.5 all cortical hem derivatives exhibited immunopositivity for βgal, including the LV choroid plexus, fimbrial glia, fimbrial neuroepithelium and some p73^+ve^ Cajal-Retzius cells within the hippocampal fissure (**Fig. 4E-H**). At E18.5, Msx1 expression was retained in the lateral ventricle choroid plexus, fimbrial neuroepithelium and some fimbrial glia (**Fig. 4F**). Furthermore, we observe co-expression of Blbp and Gfap with βgal at E18.5, confirming the radial glial identity of fimbrial glia and the fimbrial neuroepithelium (**Fig. 4H, fig. S14E**). Thus, Pre-NC/RPs contribute to the cortical hem lineage of the telencephalic midline at later stages than the hindbrain. To complement these histologic studies, we again employed scRNAseq of lineage traced *Msx1^CreERT2^; R26R-lsl-tdTomato* mice, induced at E12.5. We collected the developing telencephalon at E14.5 and E18.5, with expected recovery of various cell types **(fig. S15A-C, table S9)**. Traced tdTomato cells arising from the Pre-NC/RP lineage were subset and showed strong mapping of ZFTA-EPN tumor cells to cranial mesenchyme, cortical hem/choroid plexus, and Cajal-Retzius cells **(fig. S15D-G)**.

Using bulk RNA-sequencing comparison of PFA-EPN and ZFTA-EPN, multiple cortical hem lineage associated marker genes (*WNT3A, WNT7B* and *TP73*) are enriched in ZTFA-EPN (**Fig. 4I**)^40,43,44^. *Wnt3a*, a canonical marker of the cortical hem, and *Wnt7b* are highly expressed in the fimbrial neuroepithelium of E18.5 *Msx1^CreERT2^; R26R-lsl-LacZ* mice (**Fig. 4J**). Concordantly, *WNT3A* and *WNT7B* are specifically expressed in the tumor cells of ZFTA-EPN (**Fig. 4I,K**). The presence of MSX1 and expression of cortical hem markers (WNT3A, WNT7B), as well as Cajal-Retzius markers (TP73), along with extensive transcriptional mapping, demonstrates that ZFTA-EPNs mirror the Pre-NC/RP-derived telencephalic cortical hem lineage.

### Generating EPN-like tumors within the Pre-NC/RP lineage

Given that the *ZFTA-RELA* fusion gene is one of the few somatic mutations found in ependymoma, we sought to create a novel EPN tumor model by over-expressing it within the Pre-NC/RP lineage. Unfortunately, *Msx1^CreERT2^* mice develop postnatal craniofacial abnormalities, which limit post-natal survival. Instead, *Wnt1^Cre^* mice, which are commonly used to study the Pre-NC/RP compartment *in vivo,* were employed*^45^*. Although *Wnt1^Cre^* marks a broader set of dorsal hindbrain progenitors^46^, its early overlap with *Msx1* and *Cxcr4* at E9.5 enables targeting of candidate Pre-NC/RP cells (**fig. S16**). Indeed, *Wnt1^Cre^;lsl-hZFTA-RELA* fusion mice develop ependymoma-like tumors of the brainstem and rostral spinal cord with ∼30% penetrance. These tumors histologically resemble human ependymomas, and express ependymoma marker genes including Msx1, Gfap, and S100 (**Fig. 5A-B**). As *Wnt1^Cre^* is not expressed in the telencephalon, we expectedly did not observe any tumors in the cerebral hemispheres^47^. Although the majority of ZFTA-RELA ependymomas are found in the forebrain, primary ZFTA-RELA ependymomas have been repeatedly observed in both the posterior fossa and spinal cord^48^. We therefore interpret tumor phenotypes within the context of this shared early lineage rather than as arising from later Wnt1-derived cell types.

**Fig. 5.**
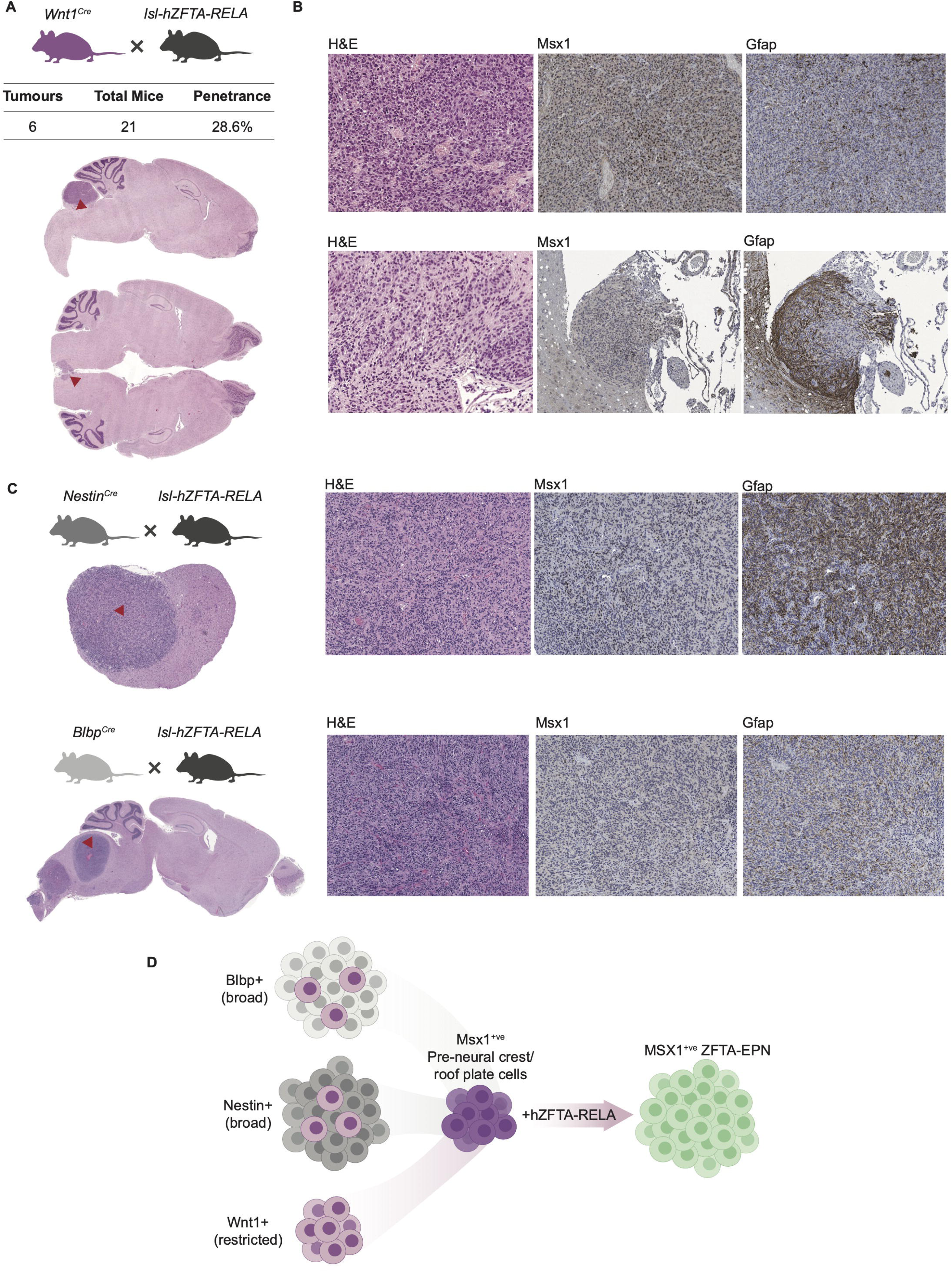
Expression of *hZFTA-RELA* in the Pre-NC/RP lineage generates EPN-like tumors. A. Pre-NC/RP lineage restricted mice (*Wnt1^Cre^*) over-expressing hZFTA-RELA generate ependymoma like tumors. Sagittal sections of whole brain stained with hematoxylin and eosin (H&E). Red arrowheads identify the lesion. B. Pre-NC/RP derived ZFTA-RELA tumors exhibit ependymoma markers. Immunohistochemical staining of sagittal sections from two *Wnt1^Cre^;lsl-hZFTA-RELA* mice showing high-magnification stains for H&E, Msx1, Gfap, and S100. C. Previously published murine models of ZFTA-EPN (*Nestin^Cre^;lsl-hZFTA-RELA, Blbp^Cre^;lsl-hZFTA-RELA)* develop ependymoma like tumors that express MSX1. Immunohistochemical staining of coronal sections showing low-magnification H&E stain (top) and high-magnification stains for H&E, Msx1, Gfap, and S100 (bottom). Red arrowheads identify the lesion. D. Proposed model for the convergence of ZFTA fusion ependymoma on Pre-NC/RP cells.

Next, we compared our model to existing mouse models of ZFTA-RELA ependymoma, by generating Cre recombinase lines targeting the radial glial lineage (Blbp-Cre or Nestin-Cre mice)^49–51^. Expectedly, these tumors exhibited ependymoma-like histology, and express ependymoma marker genes (Gfap and S100). However, excitingly they also express the Pre-NC/RP marker gene MSX1 (**Fig. 5C**). Our findings constitute the first evidence that the Pre-NC/RP lineage has the capacity to produce MSX1+ve ependymoma-like tumours *in vivo*. Furthermore, the convergence of several ZFTA-EPN models on MSX1 expression corroborates a Pre-NC/RP model of ependymoma cellular origins **(Fig. 5D)**.

## Discussion

Ependymomas transcriptionally mirror local derivatives of Pre-NC/RP cells along the dorsal neural tube, with ubiquitous MSX1 expression across all subgroups. Fascinatingly, our results suggest that PFA-EPN likely arises in the first trimester of pregnancy, when these cells transiently exist in normal development. The onset of tumor initiation during the vulnerable first weeks of fetal development as the neural tube is closing suggests that a sober second look at causative metabolic and environmental insults is required. Indeed, hypoxia has been reported as essential for normal neural tube morphogenesis in rodents, and our group previously demonstrated a similar dependence on hypoxia as a defining feature of PFA-EPN^22,52–54^. Prior epidemiological attempts to study environmental exposures in ependymoma patients have identified trends associated with exposure to carbon monoxide, nitrogen oxides, and certain pesticides^55–57^. However, the identification of relevant exposures was limited by focusing on late or post-natal exposures in a subgroup agnostic manner. We suggest that future epidemiological investigations focusing on PFA-EPN and including exposures from the first weeks of pregnancy might identify non-genetic drivers.

Our biggest impacts on health outcomes in oncology have consistently stemmed from early recognition and prevention of cancer, rather than novel treatments for advanced metastatic disease. Recognition of transformed Pre-NC/RP cells, or oncofetal antigens shed by Pre-NC/RP cells *in utero* could enable early diagnosis and treatment. Viable biomarkers to screen populations of infants for early PFA could include Pre-NC/RP specific genes not expressed after the first trimester, or low-pass detection of aneuploidy in circulating tumor DNA.

Distinctions in the incidence, age of onset, and anatomic location have long been reported between PFA and PFB ependymoma, but the basis of these differences has remained cryptogenic^1,24,29,30,58,59^. Our findings that PFA-EPNs mirror more lateral, primitive Pre-NC/RP cells in early development compared to PFB-EPNs, which resemble midline roof plate-derived radial glia from later in gestation, provide a proximate explanation for the higher incidence and early age of onset in PFA-EPN, due to the number of cells at risk for transformation. Pre-NC/RP cells also possess the capacity for significant migration, correlating with the degree of tumoral extension laterally and caudally which is frequently observed in PFA-EPN^60^. This is distinct from the small pool of RPRG cells which are anatomically restricted to the dorsal midline and differentiate later in development, corresponding to the lower incidence, midline location, and presentation in adolescents and young adults observed in PFB-EPN^30,61,62^. The fact that many of these phenotypes can be explained by considering differences in the local Pre-NC/RP derived populations reinforces the importance of investigating CNS tumorigenesis in the context of normal development.

We demonstrate for the first time that the vomiting center of the brain is partly derived from Pre-NC/RP cells. The origin of 4^th^ ventricular subependymoma from the area postrema suggests that additional rare brain tumor entities, derived from Pre-NC/RP remnant populations such as the sub-fornical and sub-commissural organs, that are not currently recognized as ependymoma variants may yet be identified^35,63^. Notably, tumors of the choroid plexus also originate from the roof plate lineage but likely arise from malignant transformation of cell types further down the differentiation hierarchy, distinguishing them from ependymomas^64^. That choroid plexus carcinomas do not evolve into ependymomas over time is further evidence against a de-differentiation model. Our findings show that ependymomas are not glial tumors derived from the neural tube but are best considered a family of ‘roof plate’ derived tumors, along with choroid plexus lesions. Indeed, the ciliated cells observed in ependymomas for many decades, presumed to be ciliated ependymal cells, are likely partially differentiated choroid plexus cells.

Intriguingly, PFA and ZFTA ependymomas make up the majority of cases. The roof plate in the telencephalon and hindbrain are greatly expanded compared to other regions of the CNS, suggesting that roof plate complexity may predispose to transformation. In the hindbrain, the expanded rhomboid roof plate is attributed to the rapid growth of the human upper and lower rhombic lips^65^. We previously published that PFA ependymoma has a dependence on hypoxic environments^22^, and propose that this may reflect the challenging and metabolically voracious microenvironment of the lateral hindbrain roof plate^22^. Whether this zone of relative starvation is responsible for the ‘hypoxophillic’ phenotype of PFA-EPN, or if it contributes to its pathogenesis will require additional experimentation.

Standard of care therapy for ependymoma remains limited to surgery and radiation, making the identification of more effective and less toxic therapies a priority. While there are a paucity of somatic drivers in ependymoma, the validated examples (i.e., *ZFTA* fusions, *NF2* mutations) are restricted to specific anatomic compartments^24,37,50,66^. Whether this restriction is due to unique transcriptional dependencies of Pre-NC/RPs and their derivatives within each compartment, synthetic lethality in other compartments, or if the mutations are simply tolerated remains to be explored. Identification of MSX1^+ve^ cells at the apex of the cellular hierarchy across ependymomas opens the door for detailed investigation of tumorigenesis in the context of normal developmental differentiation. We hope this will further our understanding of clinical phenotypic variation between ependymoma subgroups, as well as provide routes to novel therapies based on compartment and lineage specific dependencies.

## Acknowledgements

M.D.T. is a CPRIT Scholar in Cancer Research (CPRIT - RR220051). M.D.T. is the Cyvia and Melvyn Wolff Chair of Pediatric Neuro-Oncology, Texas Children’s Cancer and Hematology Center. M.D.T. is supported by the NIH (R01NS106155, R01CA159859 and R01CA255369), NCI (2P50CA127001-16), The Pediatric Brain Tumour Foundation, The Terry Fox Research Institute, The Canadian Institutes of Health Research, The Cure Search Foundation, Matthew Larson Foundation (IronMatt), Meagan’s Walk, SWIFTY Foundation, The Victory Foundation, Ependymoma Research Foundation (ERF), STOPbraintumors Foundation, The Brain Tumour Charity, Genome Canada, Genome BC, Genome Quebec, the Ontario Research Fund, Worldwide Cancer Research, V-Foundation for Cancer Research, and the Ontario Institute for Cancer Research through funding provided by the Government of Ontario. M.D.T. was also supported by a Canadian Cancer Society Research Institute Impact grant, a Cancer Research UK Brain Tumour Award, and by a Stand Up To Cancer (SU2C) St. Baldrick’s Pediatric Dream Team Translational Research Grant (SU2C-AACR-DT1113) and SU2C Canada Cancer Stem Cell Dream Team Research Funding (SU2C-AACR-DT-19-15) provided by the Government of Canada through Genome Canada and the Canadian Institutes of Health Research, with supplementary support from the Ontario Institute for Cancer Research through funding provided by the Government of Ontario. Stand Up to Cancer is a program of the Entertainment Industry Foundation administered by the American Association for Cancer Research. M.E.B. was supported by NIH R01DE027538. The authors thank Andrew Bondoc (Manager, Brain Tumour Biobank at SickKids) and recognize the Labatt Brain Tumour Research Centre and The Michael and Amira Dan Brain Tumour Bank Network. The authors thank Ariadna Villalbí for graphic design support, SickKids Imaging Facility (Toronto, Canada) for support with microscopy work, STTARR Histopathology Core (Toronto, Canada) for assistance with mouse sample processing, and Susan Archer for technical writing support. The authors wish to acknowledge the Robert Connor Dawes Foundation for funding the EPCAM series of meetings, and accelerating the pace of ependymoma research across the globe.

## Author Contributions

M.D.T. and M.E.B. led the study. P.B., S.A.K., and R.G. designed the study, performed most of the experiments and interpreted all the results. P.B. and S.A.K. performed all *in vitro* assays, lineage tracing experiments, mouse embryonic immunohistochemistry, and analyzed all newly generated and previously published data. R.G. performed all chick embryo experiments, including imaging and analysis. A.D. and W.O. performed immunohistochemistry on human and mouse tumors. P.H., J.M., O.K. conducted *in situ* hybridization and immunohistochemistry assays on human embryonic tissue. A.J.B., J.T.J., and A.N. contributed to immunohistochemistry on human autopsy samples. N.M., O.S., S.B., J.S., J.E., E.C., R.V.O., N.A. contributed to mouse sample collection and procedures. N.vK., M.V., A.W.E., W.Ong, D.F., J.Z., A.R., G.P., L.D.H., O. Saulnier, and J.J.Y.L. contributed to pre-processing, analysis and interpretation of scRNAseq data. D.P. and C.M.R. contributed to *in vitro* experiments and analysis. N.H., H.W., H.S., J.P., and E.M. assisted with mouse husbandry and housing. R.A.S, A.B., and A.L. assisted with biobanking and sample coordination. L.M.P. and K.W.Y. contributed to interpretation of patient MRI images. X.W. generated transgenic ZFTA-RELA mice, which were used in various crosses. L.M.P., X.W., A.N., C.D., A.K., K.W.Y., V.R., L.G., D.S., Q.R.L., P.H.S., J.N.R., K.J.M., M.G., R.J.G., N.J., L.D.S., and D.W.E. assisted with data interpretation and provided expert advice for experiments. P.B., S.A.K., R.G., M.E.B., and M.D.T. prepared the figures and wrote the manuscript.

## Supplemental Figure Legends

**Fig. S1.**
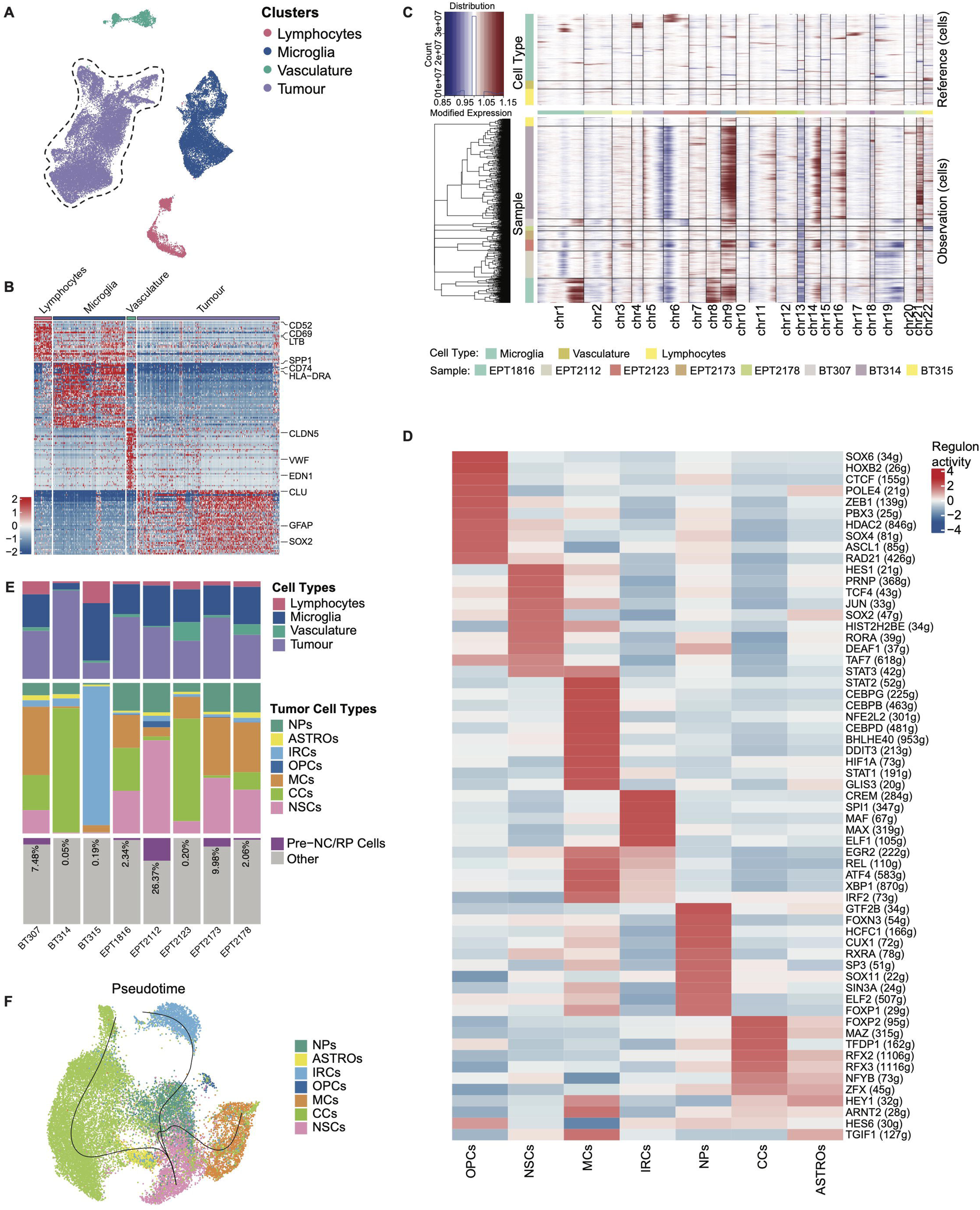
Additional mutational and transcriptional profiling of PFA-EPN scRNAseq. A. (UMAP) embedding of 41,884 PFA-EPN single-cells identifies tumor and microenvironment clusters. B. Heatmap showing relative expression of the top 50 differentially expressed genes for tumor and microenvironment clusters within PFA-EPN, highlighting 3 marker genes per cluster. Scale bar shows relative expression with a set scale -2 to 2. C. Heatmap of copy-number variations in PFA-EPN as inferred by InferCNV. Rows are grouped by tumor sample and columns represent chromosomes. D. Heatmap of the top 10 enriched transcriptional regulons as inferred by SCENIC for each tumor cell type. Scale bar shows relative regulon activity with set scale -4 to 4. E. Stacked bar plot showing the proportion of microenvironment, tumor cell types, and Pre-NC/RP annotated cells across samples. F. UMAP embedding demonstrates three unique trajectories of differentiation within PFA-EPN, as inferred from Slingshot with the NSC cluster selected as a root.

**Fig. S2.**
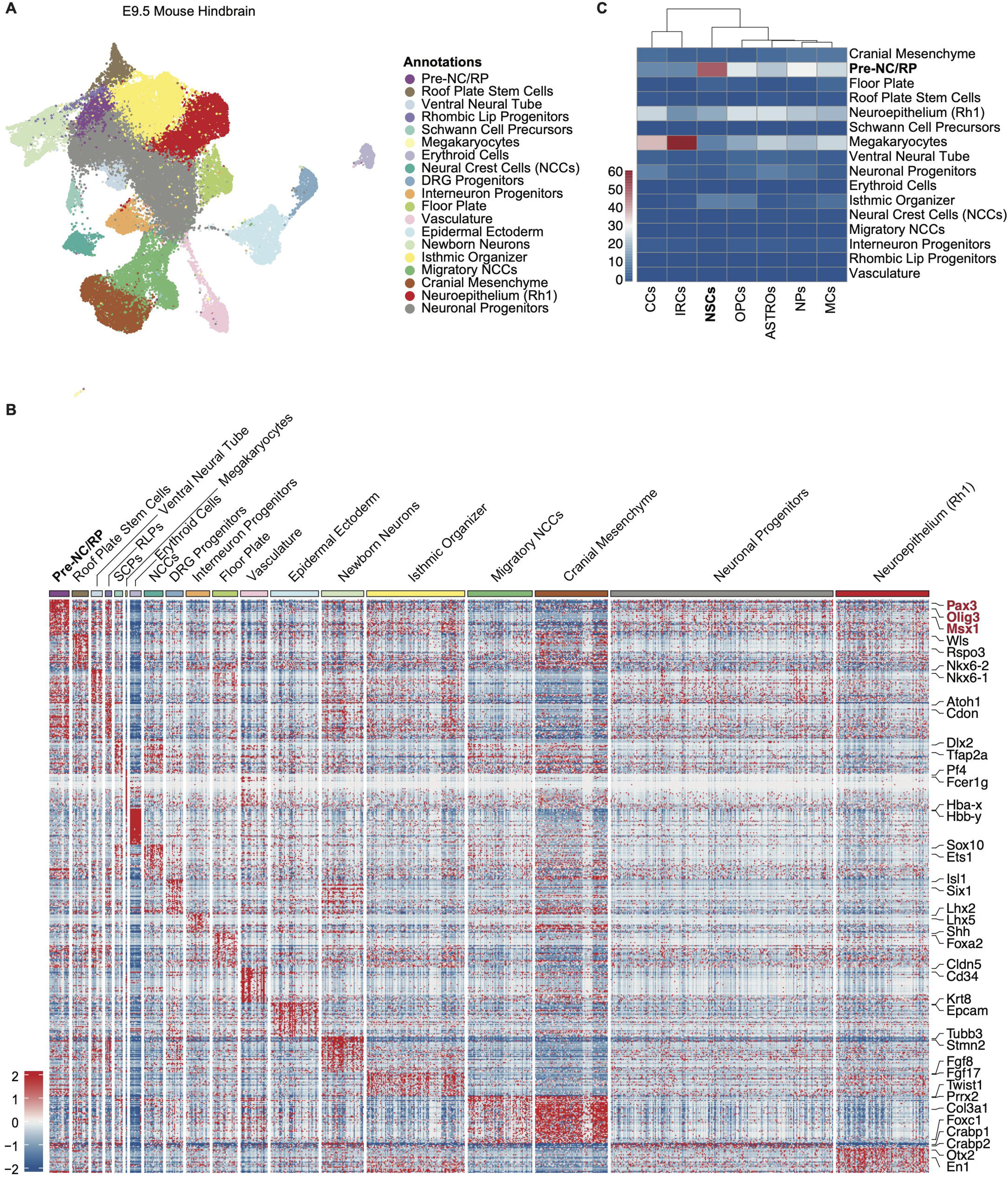
PFA-EPN resembles Pre-NC/RPs of the developing mouse hindbrain. A. UMAP embedding of the E9.5 mouse hindbrain (n=4). Clusters were manually annotated based on their unique signatures and previously published markers. B. Heatmap showing relative expression of the top 50 differentially expressed genes per cell type in the E9.5 mouse hindbrain, highlighting at least 2 genes per cell type. Set scale -2 to 2. C. Mapping of individual cells from PFA-EPN scRNAseq demonstrates resemblance to Pre-NC/RP cells of the mouse hindbrain. Heatmap shows the fraction of PFA-EPN cells that mapped to each mouse hindbrain cluster as inferred by CHETAH, by cell type annotation.

**Fig. S3.**
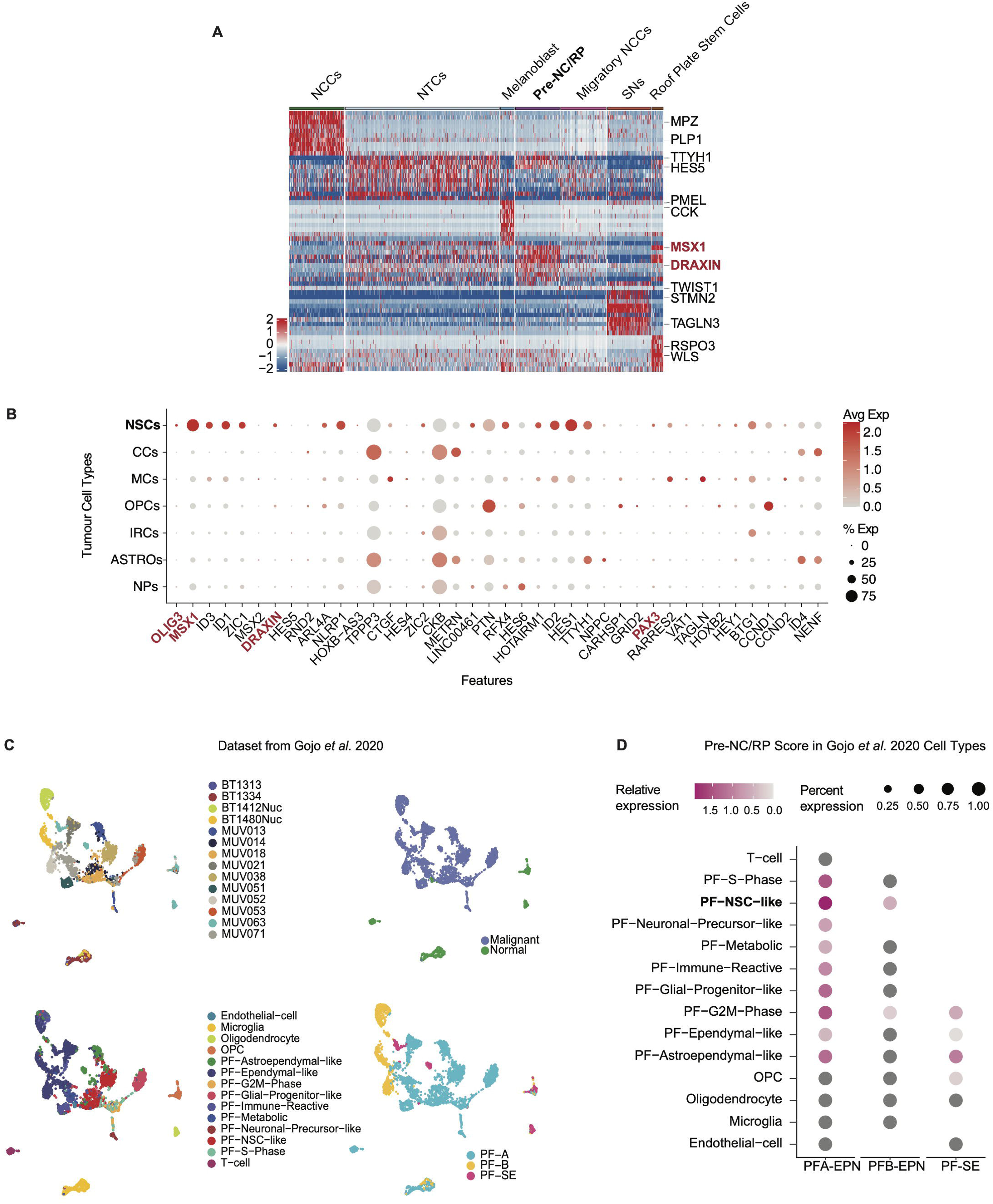
The Pre-NC/RP signature is present in posterior fossa ependymoma. A. Heatmap showing the relative expression of the top 10 differentially expressed genes per cluster from the 4 post-conception week human embryo (*Dong et al.*), highlighting at least 2 genes per cell type. Set scale -2 to 2. B. Dot plot showing expression of the top 40 genes defining Pre-NC/RPs (Dong *et al*.) in PFA-EPN by tumor cell type. Genes highlighted in red are key markers. C. Four UMAP embeddings of posterior fossa ependymoma by sample (top left), malignancy (top right), cell type (bottom left), and subgroup (bottom right). Data is re-derived from *Gojo et al.*, with annotations as previously reported. D. Dot plot showing relative expression of the Pre-NC/RP 70-gene signature in posterior fossa ependymomas (*Gojo et al.*). Set scale 0 to 2.

**Fig. S4.**
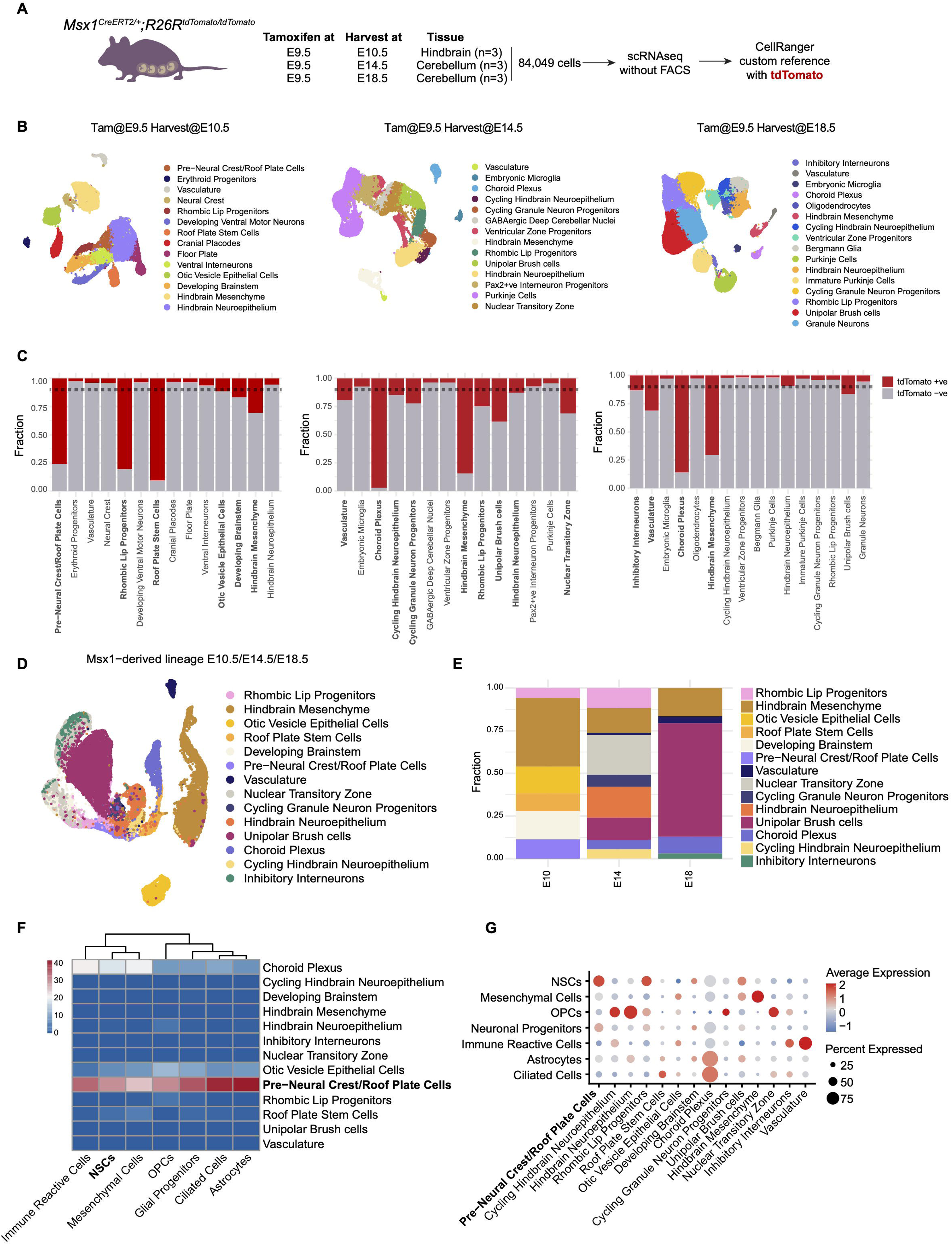
Single cell RNA sequencing of the Msx1 lineage in the developing mouse hindbrain. A. Schematic showing the workflow for scRNAseq of *Msx1Cre^ERT2/+^;R26R^Ai14/Ai14^* lineage tracing in the hindbrain. B. UMAP embedding of mouse hindbrain or cerebellum scRNAseq harvested at E10.5 (left), E14.5 (middle) and E18.5 (right) with manual annotations of cell types. C. Stacked barplots showing proportions of tdTomato^+ve^ cells (red) across cell types found in mouse hindbrain or cerebellum scRNAseq harvested at E10.5 (left), E14.5 (middle) and E18.5 (right). Dotted line represents 10% cutoff for classifying a tdTomato^+ve^ subset. D. UMAP embedding of Msx1-derived lineage in the hindbrain comprised of clusters containing more than 10% tdTomato^+ve^ cells across all timepoints. E. Stacked barplot showing proportion of Msx1-derived cell types across timepoints. F. Mapping of individual cells from PFA-EPN scRNAseq demonstrates resemblance to pre-neural crest/roof plate cells. Heatmap shows the fraction of PFA-EPN cells that mapped to each Msx1-derived lineage in the hindbrain cluster as inferred by CHETAH, by cell type annotation. G. DotPlot showing expression of gene scores in PFA-EPN tumor cells based on top-100 genes per Msx1-derived lineage cell type.

**Fig. S5.**
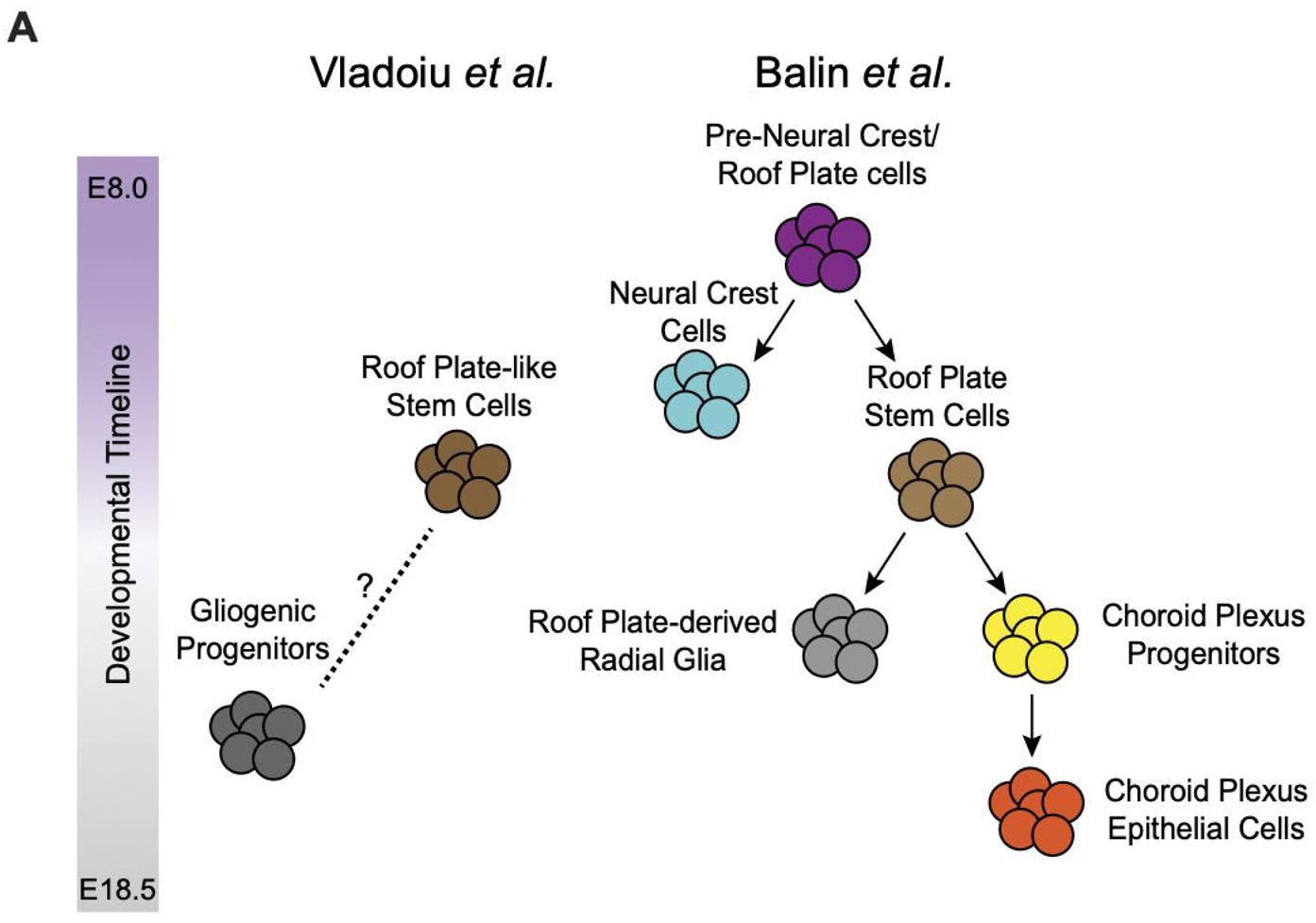
Nomenclature and relationship between putative cells of origin of ependymoma. A. An illustration showing relative hierarchies between roof plate-like stem cells and gliogenic progenitors from Vladoiu *et al*., 2019 and our reported Pre-NC/RP lineage.

**Fig. S6.**
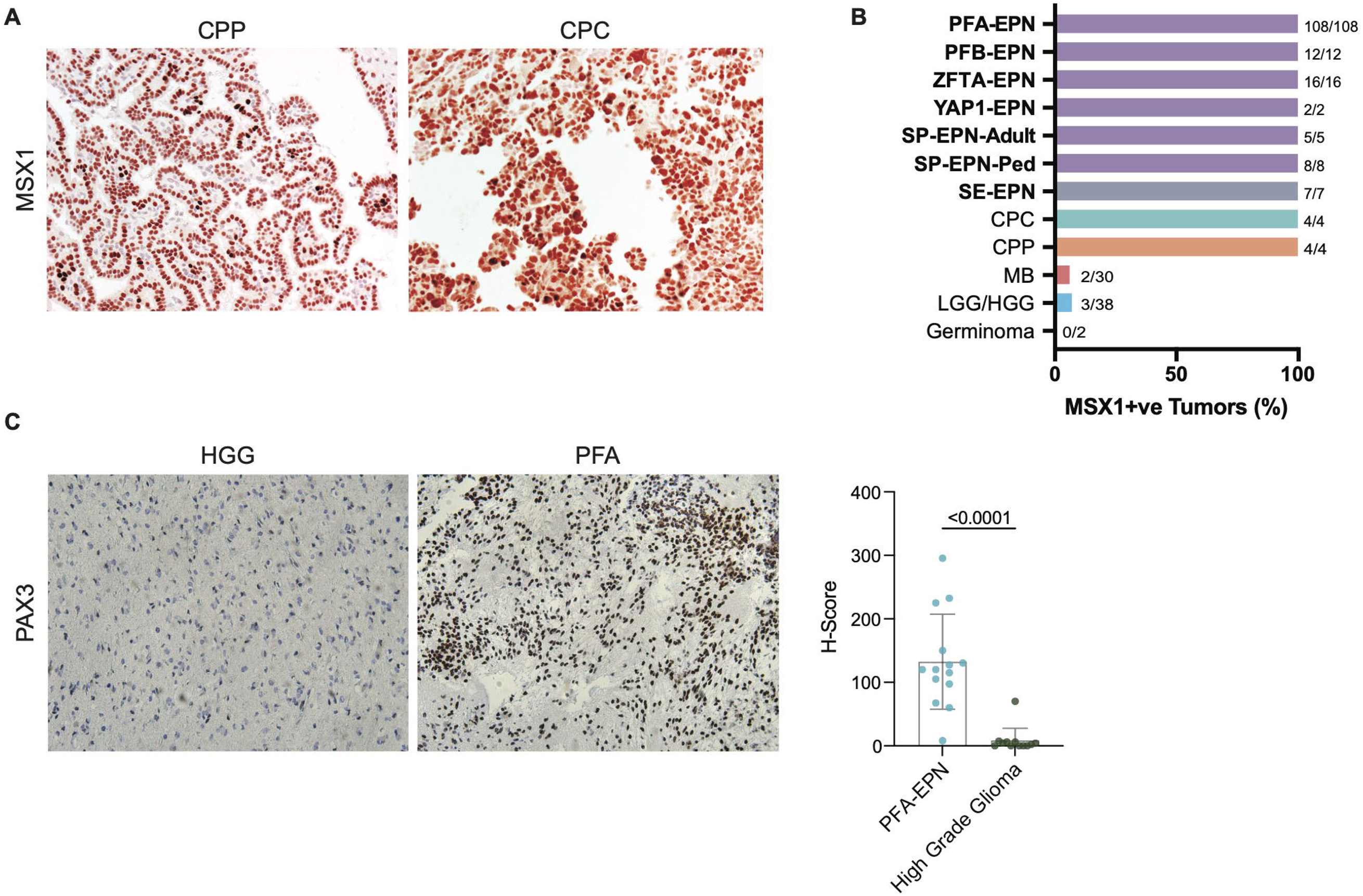
Immunohistochemical analysis of Pre-NC/RP markers in pediatric brain tumors. A. Immunohistochemistry staining shows robust MSX1 staining in choroid plexus carcinoma (CPC) and papilloma (CPP) tumors. B. Horizontal bar plot demonstrating proportion of ependymoma subgroups and control tumors that exhibit MSX1 expression by immunohistochemistry. Sample numbers noted in brackets. C. Immunohistochemical staining of the Pre-NC/RP marker PAX3 in PFA-EPN compared to high-grade gliomas (HGG). Boxplot shows median H-score with whiskers to 1.5x IQR. Statistical comparison using two-tailed unpaired T-test.

**Fig. S7.**
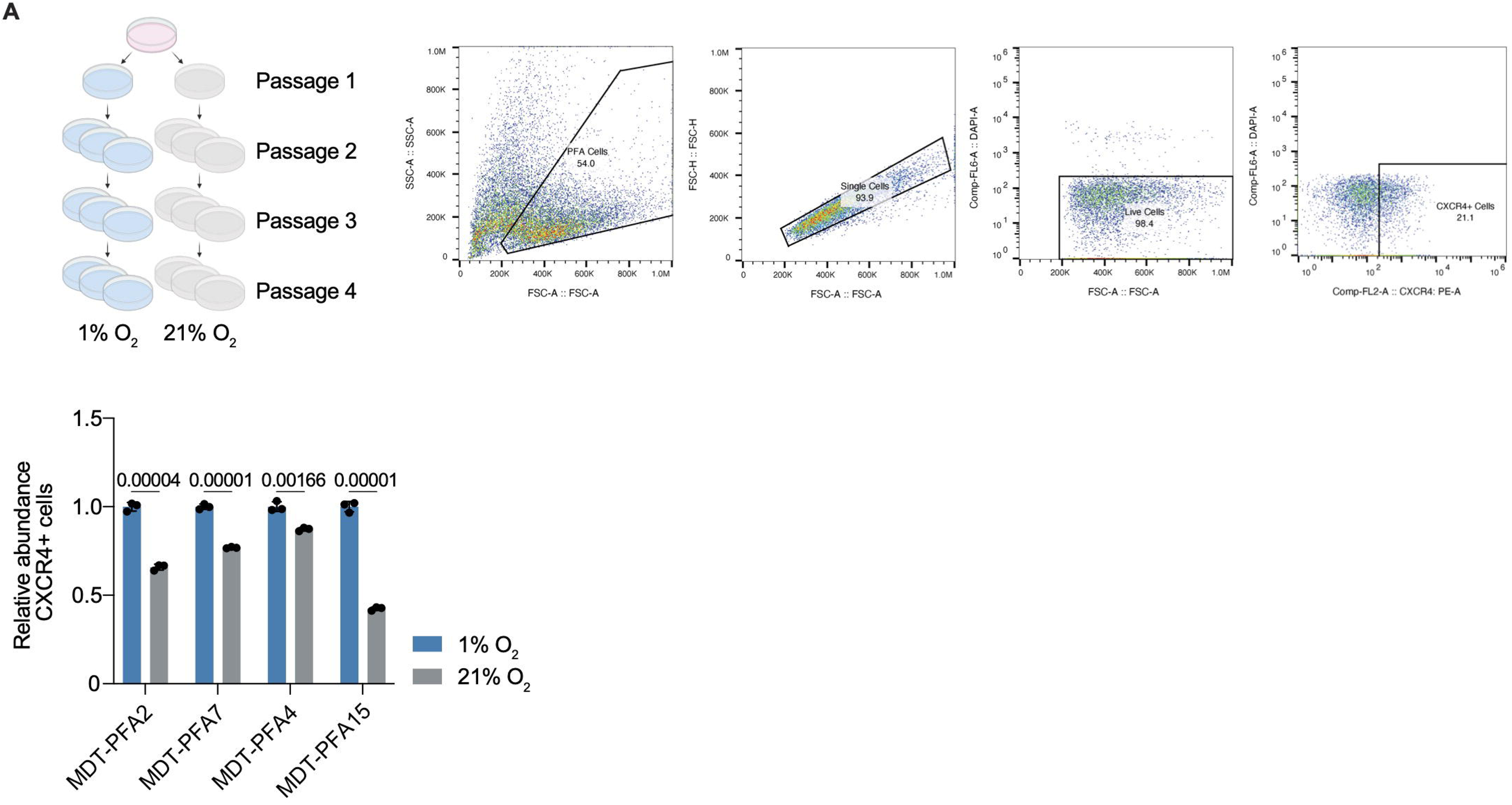
Analysis of stem cell abundance in PFA-EPN in varying oxygen conditions. A. PFA-EPN primary tumor cells cultured in hypoxia (1% O2) have a higher abundance of CXCR4+ve stem cells compared to normoxia (21% O2). Diagram illustrating the experimental culture set up, representative gating strategy, and bar plot showing relative abundance of CXCR4^+ve^ stem cells as determined by flow cytometry. Statistical comparison using two-tailed unpaired T-test.

**Fig. S8.**
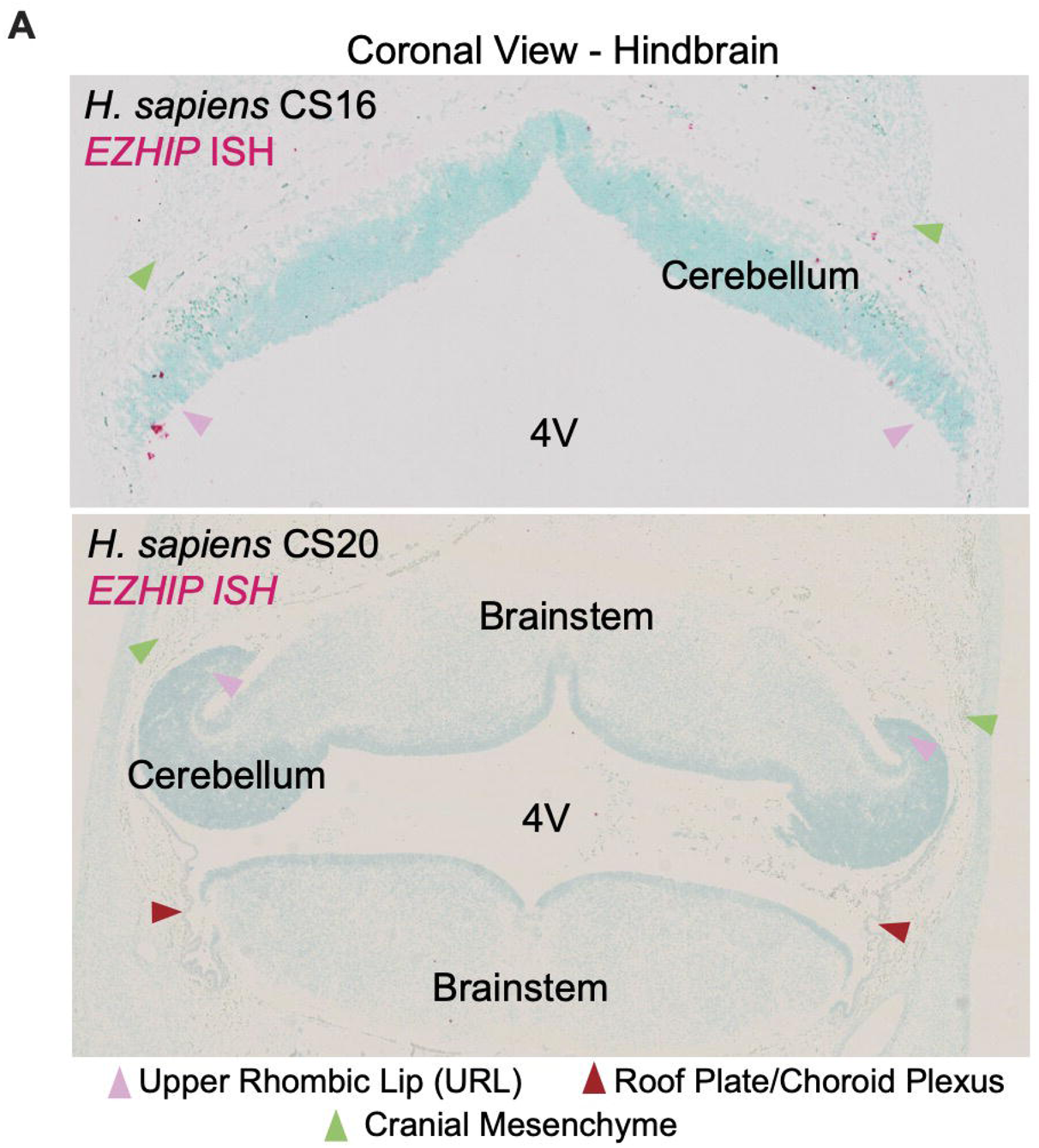
Analysis of EZHIP expression in embryonic human hindbrain. A. *In situ* hybridization expression of *EZHIP* in coronal sections of the developing human hindbrain at CS16 (top) and CS20 (bottom). Arrowheads denote anatomical location.

**Fig. S9.**
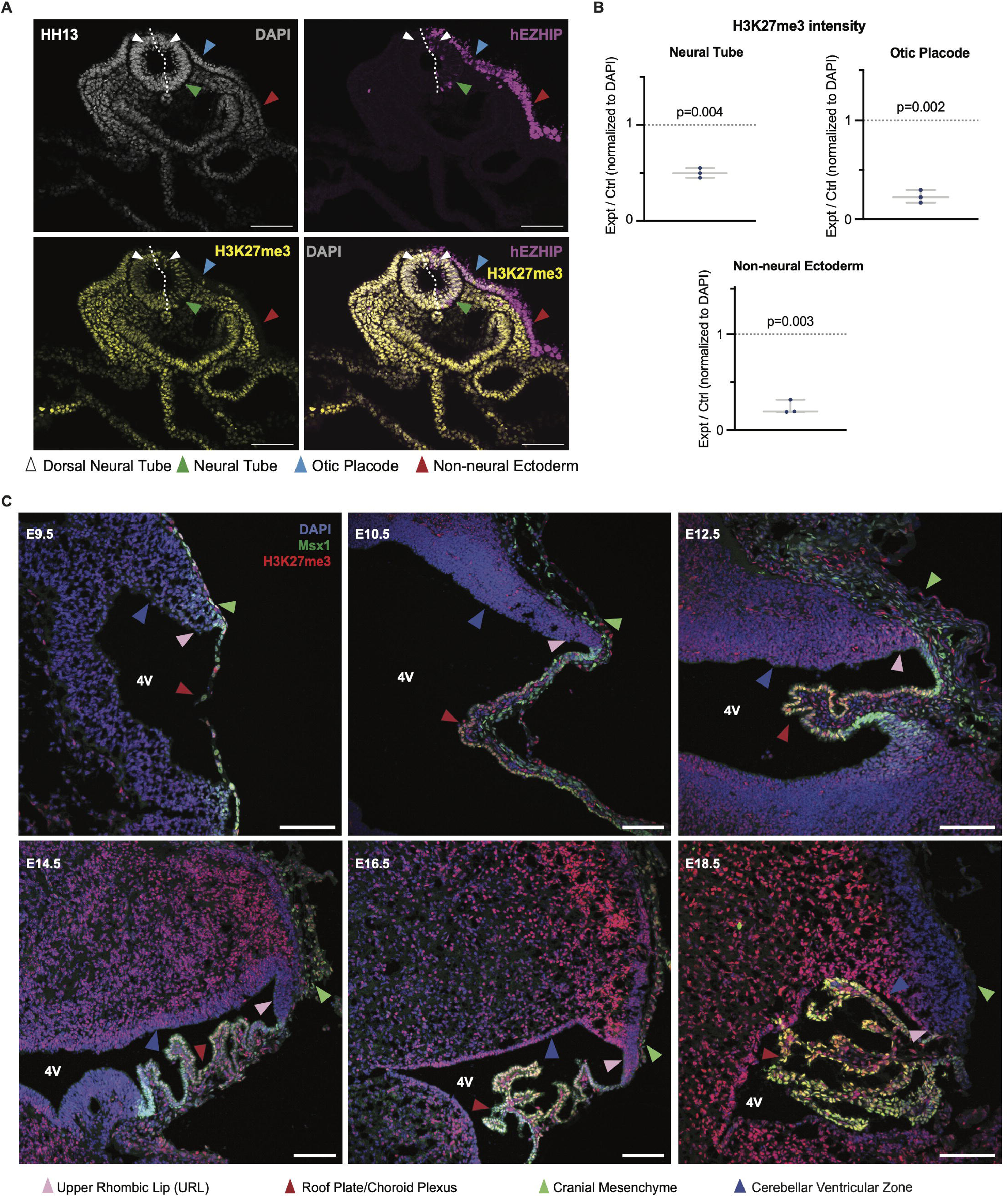
Expression of histone H3 lysine-27 trimethylation (H3K27me3) in chick and mouse embryos. A. Immunohistochemical staining of axial sections through chick neural tube at HH13, stained for DAPI (grey), hEZHIP (purple), H3K27me3 (yellow, top). Scale bar 100μm. Arrowheads denote anatomical location. B. Boxplots showing quantification of relative H3K27me3 intensity within the neural tube (left), otic placode (right) and non-neural ectoderm (bottom). Arrowheads indicate anatomic structure. Statistical comparison using a two-tailed paired T-test. Centre line shows median and lower and upper whiskers extend 1.5× the IQR. C. H3K27me3 marks are deposited to the roof plate and choroid plexus as development progresses. Immunofluorescence of sagittal sections through mouse embryonic hindbrains across development (E9.5 – E18.5) stained for Msx1 (green), H3K27me3 (red) and DAPI (blue). Scale bar 100μm.

**Fig. S10.**
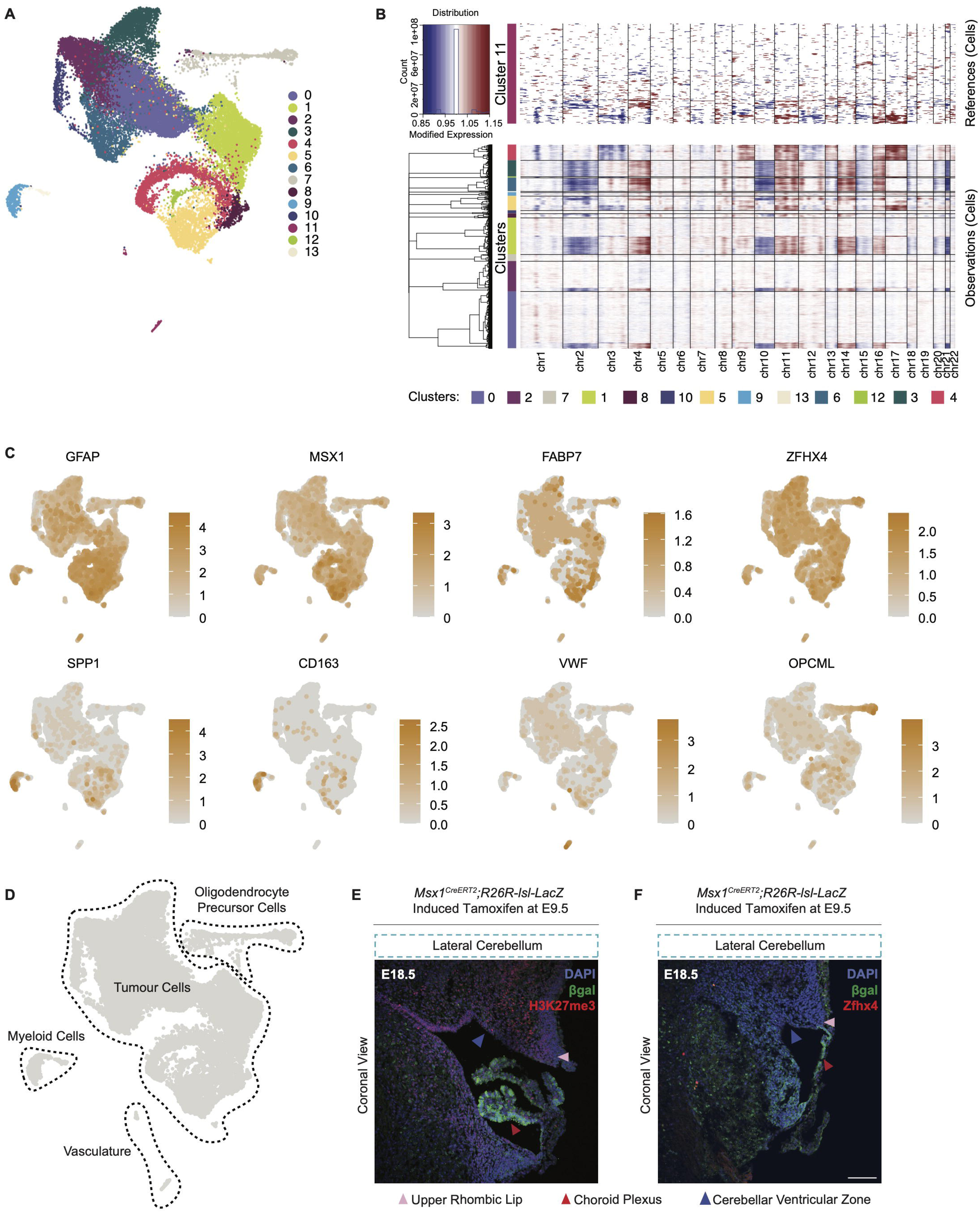
Characterization of PFB-EPN single nucleus RNA sequencing. A. UMAP embedding of PFB-EPN tumor cells analyzed by snRNAseq, highlighting transcriptionally distinct clusters. B. Heatmap of copy-number variations in PFB-EPN as determined by InferCNV. Rows are grouped by tumor clusters and columns represent chromosomes. C. UMAP embeddings of PFB-EPN showing expression of select marker genes for tumor cells (GFAP, MSX1, FABP7, ZFHX4), microglia (SPP1, CD163), vasculature (VWF), and oligodendrocyte precursor cells (OPCML). D. UMAP embedding of PFB-EPN snRNAseq with manual annotation of tumor cells and associated micro-environment. E. Coronal section of the lateral cerebellum at E18.5 in lineage traced *Msx1^CreERT2^;R26R-lsl-LacZ* mice, stained for βgal (green), H3K27me3 (red) and DAPI (blue). Scale bar 100μm. Arrowheads denote anatomical location. F. Coronal section of the lateral cerebellum at E18.5 in lineage traced *Msx1^CreERT2^;R26R-lsl-LacZ* mice, stained for βgal (green), Zfhx4 (red) and DAPI (blue). Scale bar 100μm. Arrowheads denote anatomical location.

**Fig. S11.**
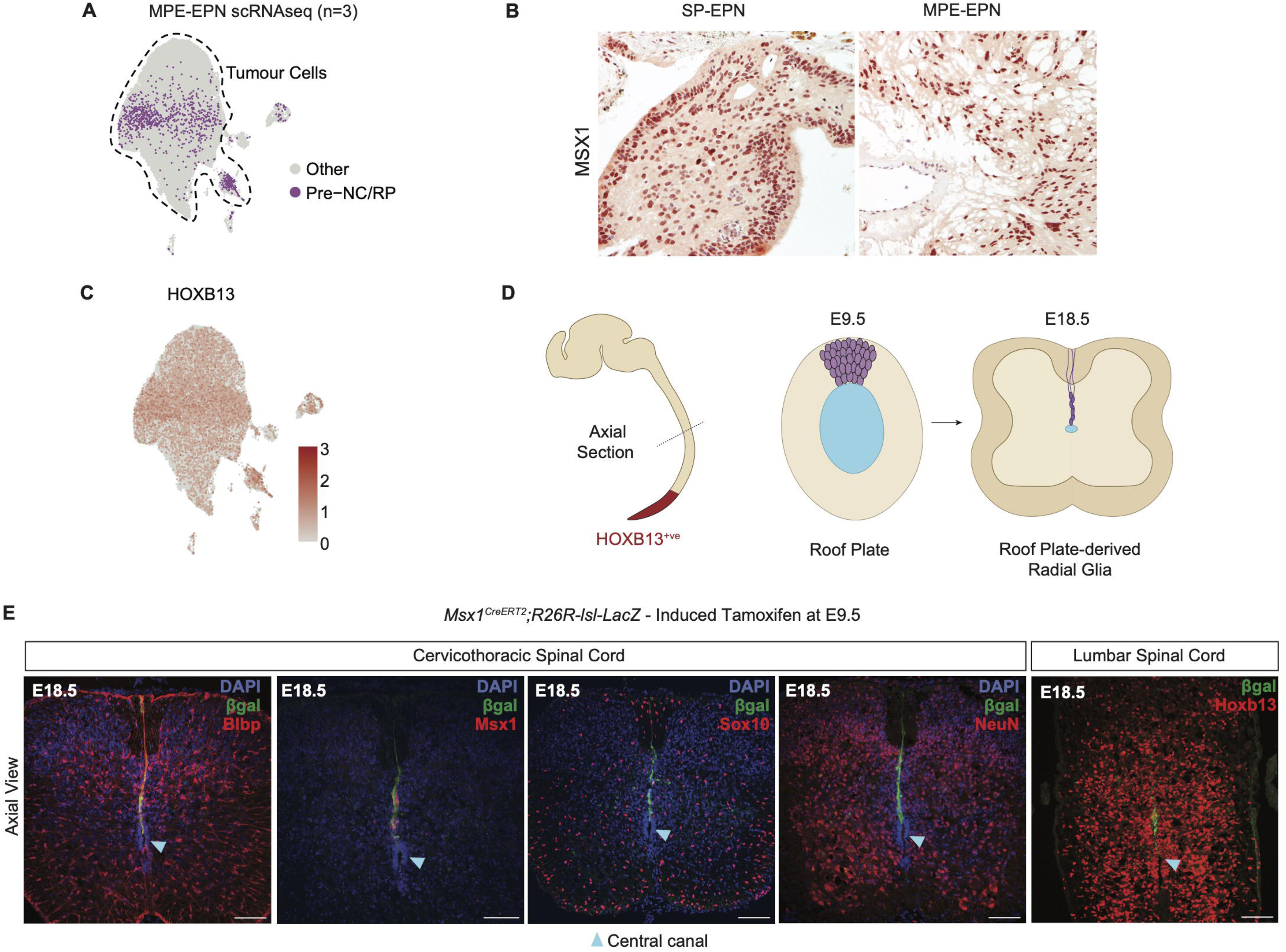
Spinal ependymoma variants resemble roof plate-derived radial glia. A. UMAP embedding of MPE-EPN tumor cells (snRNAseq), highlighting cells enriched for a Pre-NC/RP signature. B. Immunohistochemical staining of MSX1 in patient samples is positive (red) in all SP-EPN and MPE-EPN tumors (n=13). C. UMAP embedding of MPE-EPN demonstrates expression of *HOXB13* within tumor cells. D. Illustration of axial sections through the E9.5 (left) and E18.5 (right) murine spinal cord, highlighting morphological transformation of roof plate cells to roof plate-derived radial glia. E. Pre-NC/RPs give rise to roof plate-derived radial glia, but not oligodendrocyte (Sox10) or neuronal (NeuN) populations. Axial sections through the cervicothoracic (left) and lumbar (right) spinal cord at E18.5 in lineage traced *Msx1^CreERT2^;R26R-lsl-LacZ* mice induced with tamoxifen at E9.5, stained for βgal (green), DAPI (blue), and Blbp, Msx1, Sox10, NeuN, Hoxb13 (red, left to right). Scale bar 100μm. Arrowhead denotes central canal of spinal cord.

**Fig. S12.**
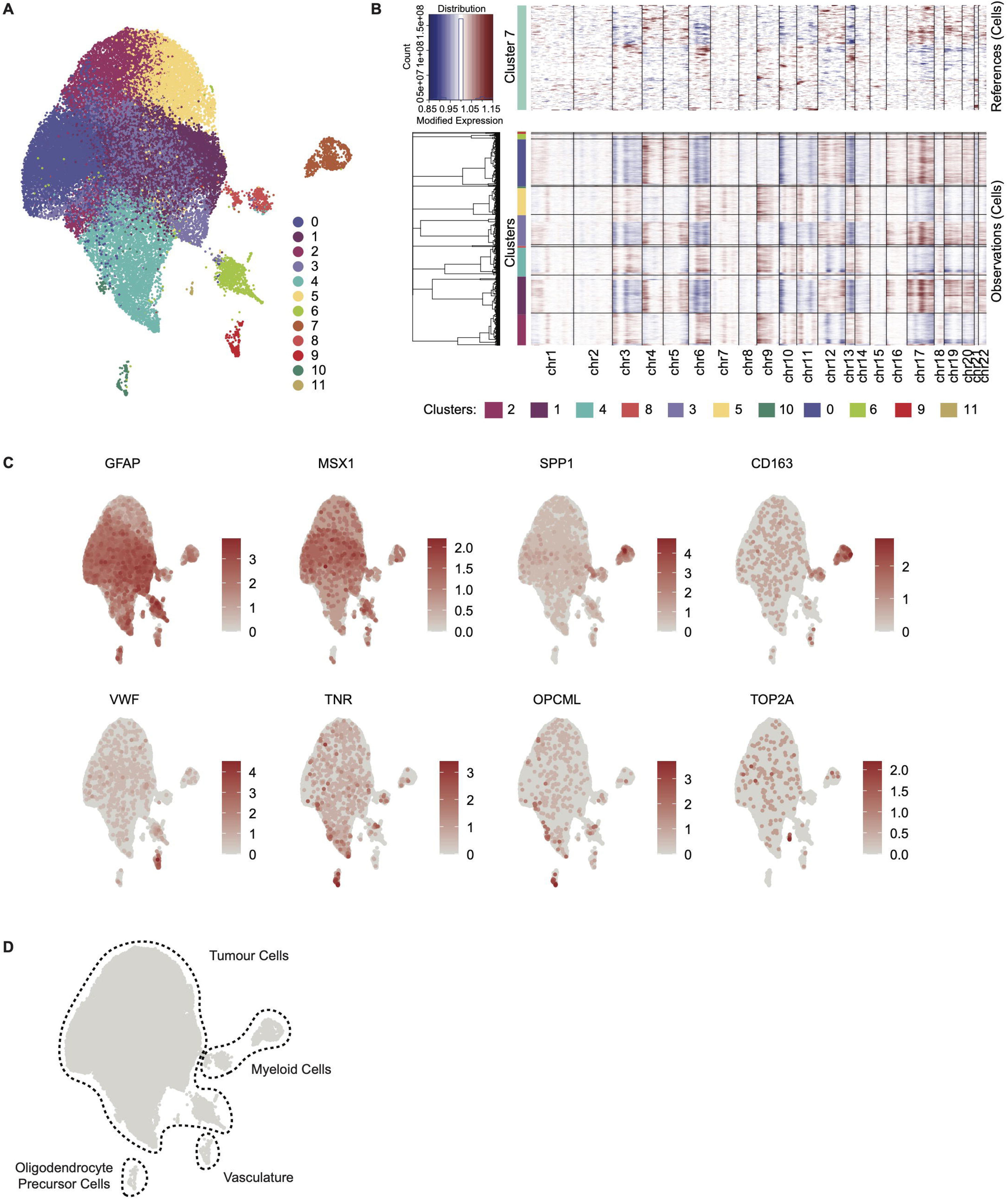
Characterization of MPE-EPN single nucleus RNA sequencing. A. UMAP embedding of MPE-EPN tumor cells analyzed by snRNAseq, highlighting transcriptionally distinct clusters. B. Heatmap of copy-number variations in MPE-EPN as determined by InferCNV. Rows are grouped by tumor clusters and columns represent chromosomes. C. UMAP embeddings of MPE-EPN showing expression of select marker genes for tumor cells (GFAP, MSX1), microglia (SPP1, CD163), vasculature (VWF), oligodendrocyte precursor cells (OPCML, TNR) and cycling cells (TOP2A). D. UMAP embedding of MPE-EPN snRNAseq with manual annotation of tumor cells and associated micro-environment.

**Fig. S13.**
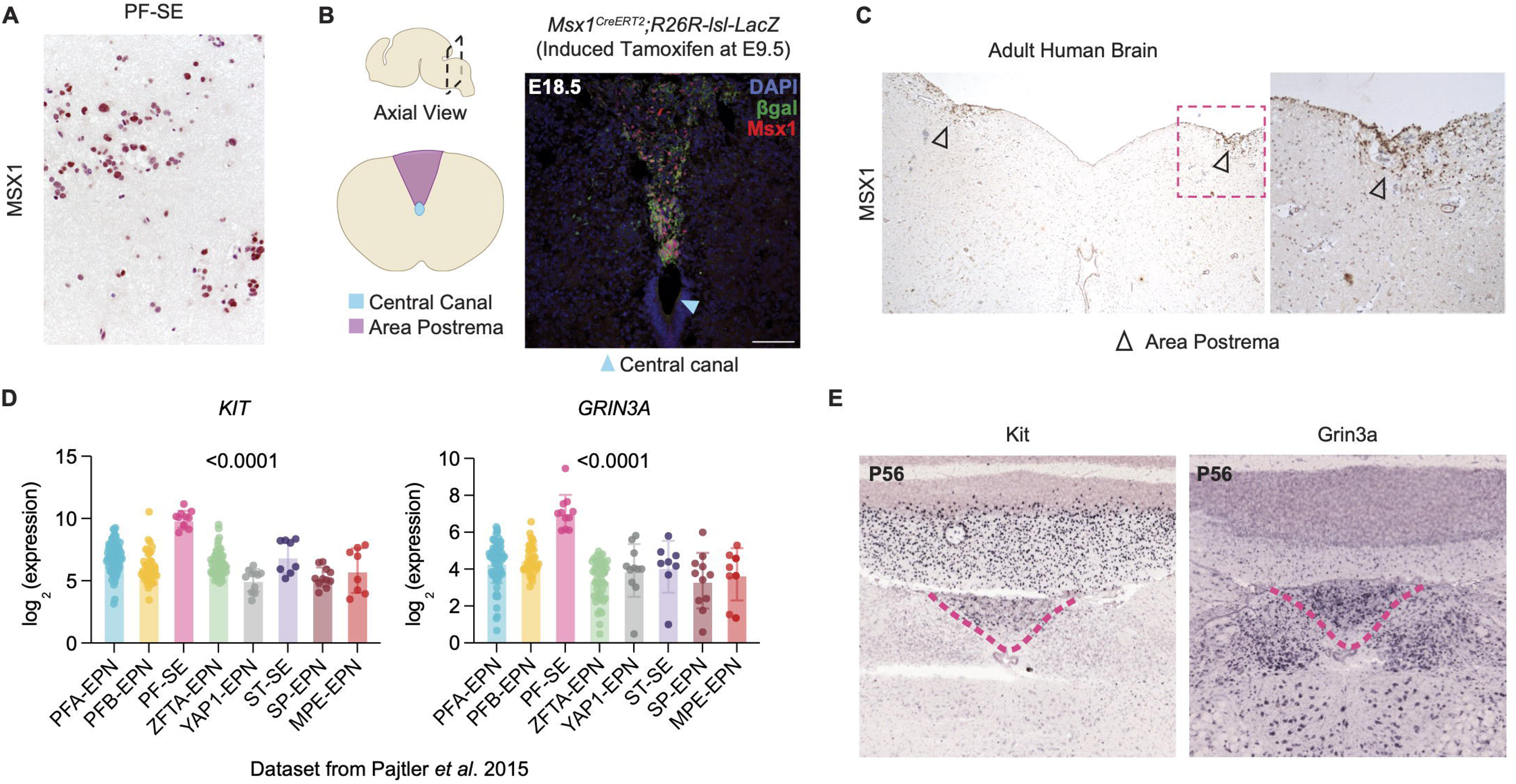
Posterior fossa subependymoma mirrors dorsal midline-derived area postrema. A. Immunohistochemical staining of MSX1 in patient samples is positive (red) in all subependymoma tumors (n=7). PF-SE denotes posterior fossa subependymoma. B. Pre-NC/RP derivatives give rise to the area postrema. Illustration (left) of a coronal section through the caudal hindbrain highlighting the anatomical location of the area postrema. Coronal section (right) through the area postrema at E18.5 in lineage traced *Msx1^CreERT2^;R26R-lsl-LacZ* mice induced with tamoxifen at E9.5, stained for βgal (green), Msx1 (red) and DAPI (blue). Scale bar 100μm. C. Immunohistochemical staining of MSX1 in autopsy samples of the adult human caudal hindbrain is positive (brown) in the paired area postrema (arrowheads). Inset (right) shows higher magnification. D. Bar plot showing expression of PF-SE marker genes *KIT* and *GRIN3A* across nine ependymoma subgroups. Statistical comparison performed using two-tailed unpaired T-test. E. Representative *in situ* hybridization images from coronal sections through the area postrema in adult mice at postnatal day 54 (P54), showing expression of Kit (left) and Grin3a (right). Open-source data from the Allen Brain Atlas.

**Fig. S14.**
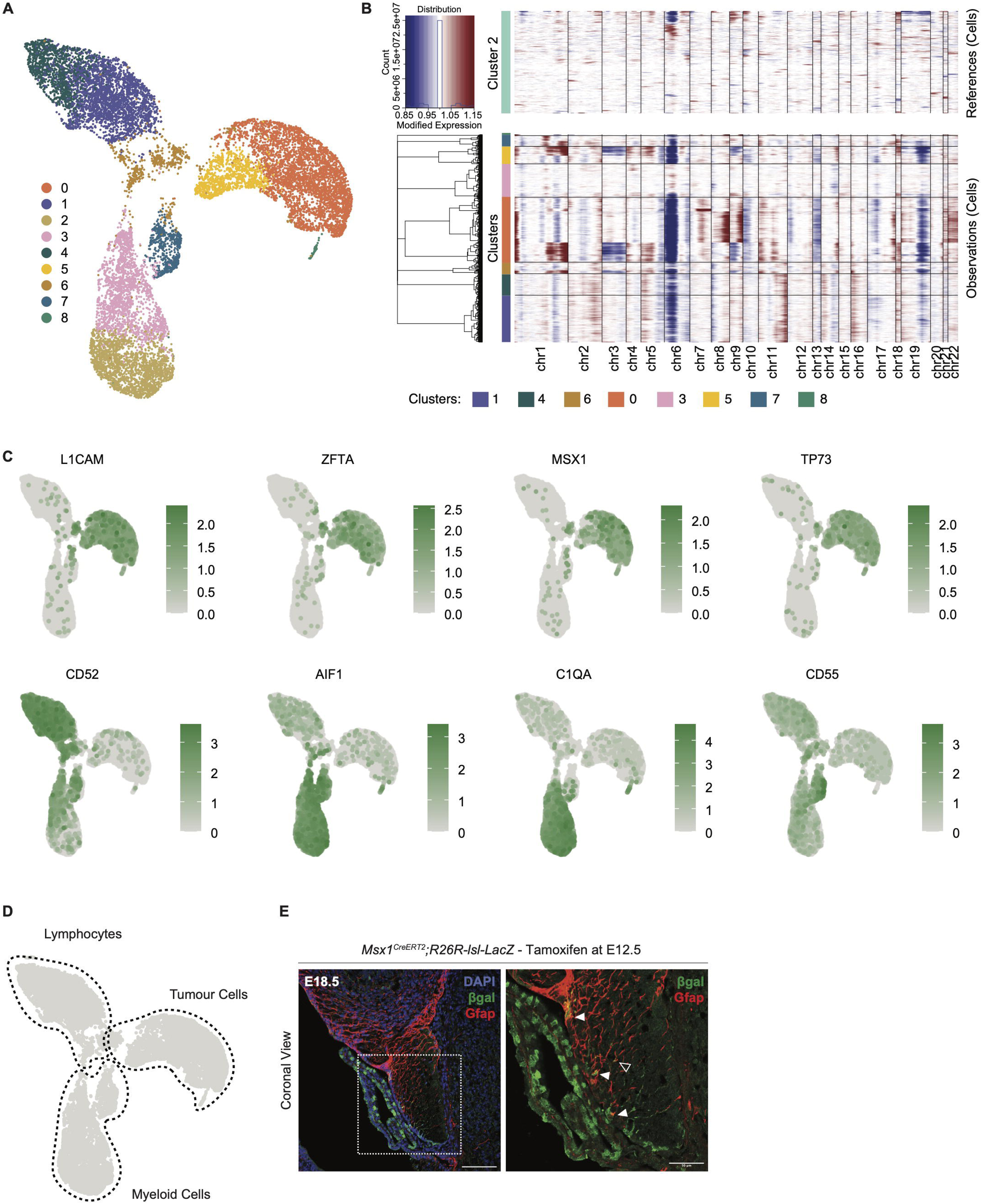
Characterization of ZFTA-EPN single cell RNA sequencing. A. UMAP embedding of ZFTA-EPN tumor cells analyzed by scRNAseq, highlighting transcriptionally identified clusters. B. Heatmap showing copy-number variations in ZFTA-EPN as deteremined by InferCNV). Rows are grouped by tumor clusters and columns represent chromosomes. C. UMAP embeddings of ZFTA-EPN demonstrating expression of select marker genes for tumor cells (L1CAM, MSX1), microglia (SPP1, CD163), vasculature (VWF), oligodendrocyte precursor cells (OPCML, TNR) and cycling cells (TOP2A). D. UMAP embedding of ZFTA-EPN scRNAseq with manual annotation of tumor cells and associated micro-environment. E. Immunofluorescence image of coronal sections through the cortical hem at E18.5 in lineage traced *Msx1^CreERT2^;R26R-lsl-LacZ* mice induced with tamoxifen at E12.5, stained for lgal (green), Gfap (red) and DAPI (blue). Inset (right) identifies βgal/Gfap double positive fimbrial glia (empty arrowheads) and fimbrial neuroepithelium (filled arrowheads). Scale bar 100μm.

**Fig. S15.**
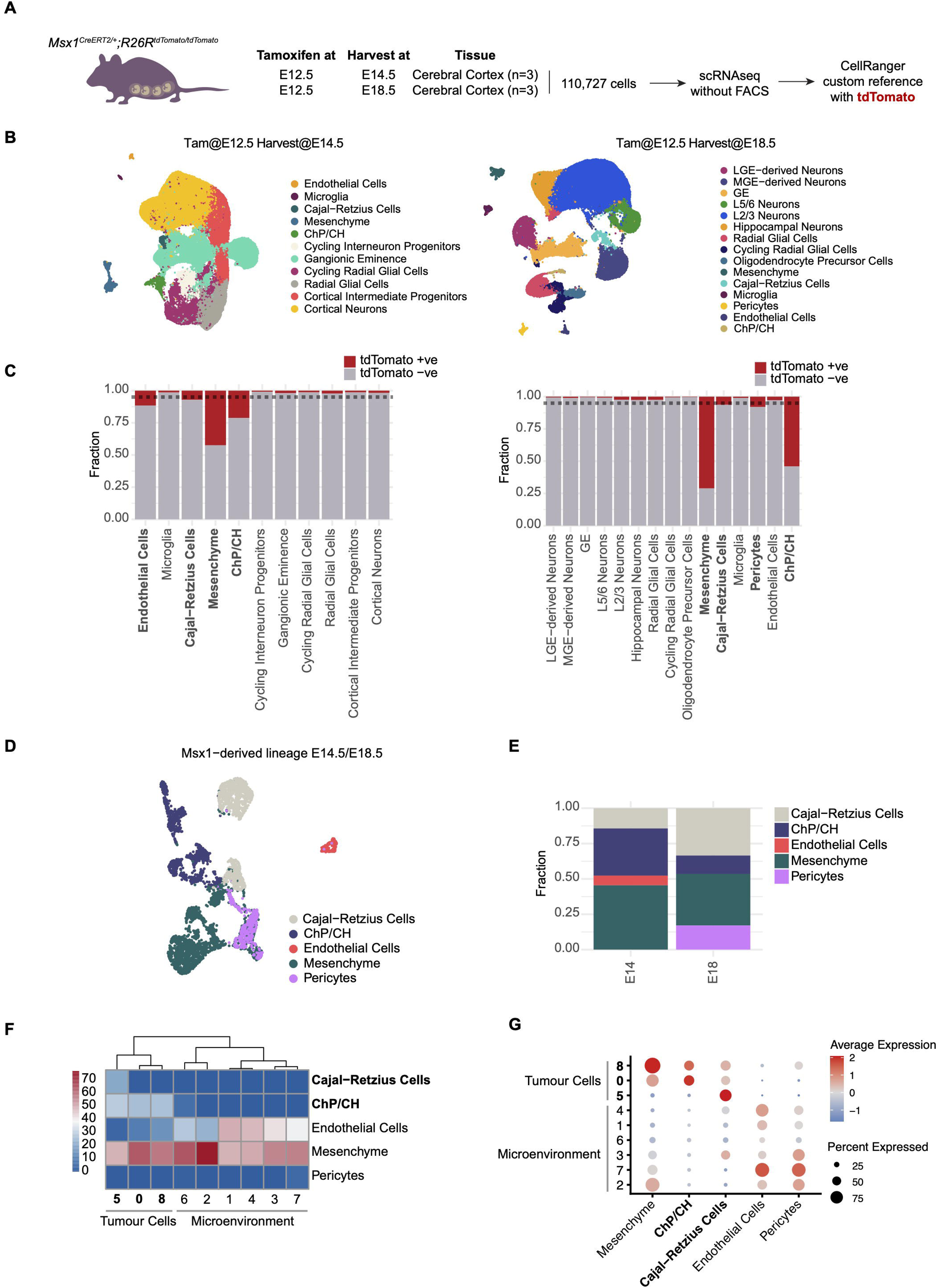
Single cell RNA sequencing of the Msx1 lineage in the developing mouse telencephalon. A. A schematic showing the workflow for scRNAseq of *Msx1Cre^ERT2/+^;R26R^Ai14/Ai14^* lineage tracing in the telencephalon. B. UMAP embedding of mouse telencephalon scRNAseq harvested at E14.5 (left) and E18.5 (right) with manual annotations of cell types. C. Stacked barplots showing proportions of tdTomato^+ve^ cells (red) across cell types found in mouse telencephalon scRNAseq harvested at E14.5 (left) and E18.5 (right). Dotted line represents 5% cutoff for classifying a tdTomato^+ve^ subset. D. UMAP embedding of Msx1-derived lineage in the telencephalon comprised of clusters containing more than 10% tdTomato^+ve^ cells across all timepoints. E. Stacked barplot showing proportion of Msx1-derived cell types across timepoints. F. Mapping of individual cells from ZFTA-EPN scRNAseq demonstrates resemblance of tumor cells ChP/CH and Cajal-Retzius cells. Heatmap shows the fraction of ZFTA-EPN cells that mapped to each Msx1-derived lineage in the telencephalon cluster as inferred by CHETAH, by cell type annotation. G. DotPlot showing expression of gene scores in ZFTA-EPN tumor cells based on top-100 genes per Msx1-derived lineage cell type.

**Fig. S16.**
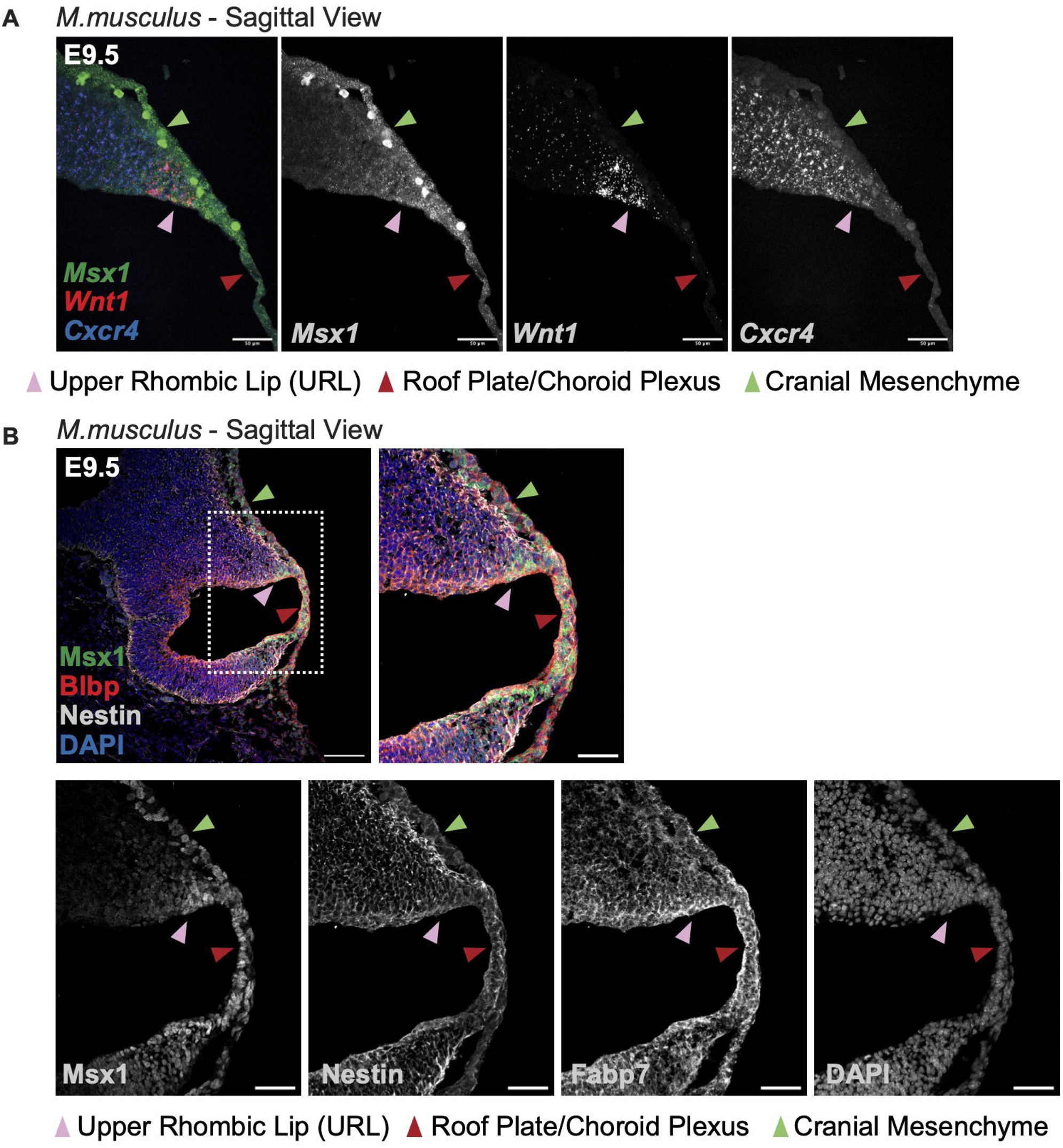
Expression of progenitor markers in the developing mouse upper rhombic lip. A. Multiplexed RNAscope *in situ* hybridization of the E9.5 developing mouse hindbrain, showing co-expression of *Msx1* (green), *Wnt1* (red), and *Cxcr4* (blue) in the upper rhombic lip. Individual black and white channels for each probe are shown for clarity. Arrowheads indicate anatomic structures. Scale bar 50μm. B. Immunofluorescence through sagittal sections of the developing mouse hindbrain at E9.5 shows overlapped expression between Msx1 (green), Blbp (red) and Nestin (gray) in the upper rhombic lip. Scale bar is 100μm. Individual black and white channels for each probe are shown for clarity. Arrowheads indicate anatomic structures. Scale bar 50μm.

## Supplemental Data Tables

**Table S1 – Tumor cluster-specific gene expression in PFA-EPN, related to Fig. 1**.

**Table S2 – Cluster-specific gene expression in E9.5 mouse hindbrain scRNAseq, related to fig. S2.**

**Table S3 – Gene signature for Pre-NC/RP lineage as determined by human fetal scRNAseq, related to Fig. 1, 3-5.**

**Table S4 – Cluster-specific gene expression across *Msx1^CreERT2^;R26R^Ai14^* hindbrains related to fig. S4.**

**Table S5 – Markers expressed in roof plate-like stem cells and gliogenic progenitors from Vladoiu *et al*., 2019 and the Pre-NC/RP lineage derivatives related to fig. S5.**

**Table S6 – Cluster-specific gene expression in PFB-EPN, related to Fig. 3, fig. S10.**

**Table S7 – Cluster-specific gene expression in MPE-EPN, related to fig. S11-12.**

**Table S8 – Cluster-specific gene expression in ZFTA-EPN, related to Fig. 4, fig. S14.**

**Table S9 – Cluster-specific gene expression across *Msx1^CreERT2^;R26R^Ai14^* forebrains related to fig. S15.**

**Table S10 – Demographics for all single cell/nucleus samples, related to Fig. 1,3-5 and fig. S4-15.**

## Methods

### Human tumor collection

Patient tumor samples used in this study were obtained with informed consent of patients. All experimental procedures with collected tissues were performed in compliance with Research Ethic Board at The Hospital for Sick Children (REB 1000055059) and Cincinnati Children’s Hospital Medical Centre. All human cerebellar samples utilized in this study were procured under protocols approved by the Institutional Review Board (IRB) of Seattle Children’s Research Institute. Samples were sourced from the Human Developmental Biology Resource (HDBR) at University College London and Newcastle University in the United Kingdom, where each sample was gathered with prior patient consent and in full compliance with institutional and legal ethical standards. FFPE samples of autopsy brain from 3 older adult individuals were obtained from the Calgary Brain Bank. Sections of the following areas were examined: pons, medulla, spinal cord (representative from all levels), vermis, and caudate head (with adjacent lateral ventricle). Ethics approval was provided by the Conjoint Health Research Ethics Board (CHREB) of the University of Calgary and the Health Research Ethics Board of Alberta (HREBA).

### Magnetic Resonance Imaging

All MRI scans were performed on patients with ependymoma from the Hospital for Sick Children, Toronto, Canada.

### Mouse husbandry and handling

All mouse handling was performed in compliance with standard operating procedures of The Centre for Phenogenomics (TCP, Toronto) and by the Institutional Animal Care and Use Committee of the Center for Comparative Medicine (CCM) at Baylor College of Medicine. For embryo collection, female mice were placed on time-controlled matings, with the day of the plug defined as E0.5. The following commercial wildtype strains were purchased from Jackson Laboratory: *C57BL/6J* (https://www.jax.org/strain/000664), *FVB/NJ* (https://www.jax.org/strain/001800). The following mutant strains were purchased from Jackson Laboratory: *Msx1^CreERT2^*(https://www.jax.org/strain/027850), *Rosa26R^LacZ^*(https://www.jax.org/strain/003474), *Rosa26R^Ai14^* (https://www.jax.org/strain/007914),*Wnt1^Cre2^*(https://www.jax.org/strain/022501), *Nestin^Cre^*(https://www.jax.org/strain/003771). *Blbp^Cre^*mouseline was acquired from Nathaniel Heintz at the Rockefeller University. The construct built for the transgenic *lsl-hZFTA-RELA* (*lsl-hC11orf95-RELA*) mouse was CAG-NeoSTOP-flox-hZFTA-RELA-T2A-mCherry. Previously published *C11orf95-RELA* sequence was utilized^1^. *C11orf95-RELA* fusion DNA was amplified from an ependymoma cDNA pool by high fidelity Q5 polymerase (M0491S, New England Biolabs). T2A-mCherry (800bp) was amplified from pcDNA3.1(-)T2A- mCherryWPRE by high fidelity Q5 polymerase. The sequences of PCR products were validated by Sanger sequencing. Twenty-one pups were born from pronuclear injection (done with TCP Model production service). Genotyping primers target the mCherry tag.

To genotype the mutant strains, the following primers were used: *Msx1^CreERT2^* (Forward -TGA TCC AGG GCT GTC TCG; Reverse - CCC CCT CCT CAT CTG TGC; Common - CGG TTA TTC AAC TTG CAC CA), *Rosa26* WT (Forward - AAG GGA GCT GCA GTG GAG TA; Reverse - CCG AAA ATC TGT GGG AAG TC), *Rosa26R^Ai14^*(Forward - CTG TTC CTG TAC GGC ATG G; Reverse - GGC ATT AAA GCA GCG TAT CC),*Wnt1^Cre2^*(Forward - CAG CGC CGC AAC TAT AAG AG; Reverse - CAT CGA CCG GTA ATG CAG), *Rosa26R^LacZ^* (Forward - AAA GTC GCT CTG AGT TGT TAT; Reverse - GGA GCG GGA GAA ATG GAT ATG; Common - GCG AAG AGT TTG TCC TCA ACC), *LsL-ZFTA-RELA-mCherry* (Forward - CGA GGA GGA TAA CAT GGC CAT C; Reverse - CAT CAC GCG CTC CCA CTT GAA G), *BlbpCre/NestinCre* (primers for Cre, Forward - GTC GAT GCA ACG AGT GAT GAG; Reverse - CGA CGA TGA AGC ATG TTT AGC TG).

### Mouse sample collection

Mouse embryos were collected at embryonic days E9.5, E10.5, E11.5, E12.5; embryonic brains were collected at embryonic days E14.5, E16.5, E18.5 and postnatal day P56. Samples were collected dissected in cold 1xPBS and transferred to 4% paraformaldehyde for 3 hours for E9.5, E10.5; 4 hours for E11.5, E12.5, E14.5; 5 hours for E18.5 and 6 hours for P56. Samples were then washed in 1xPBS overnight at 4°C, cryoprotected in 30% sucrose at 4°C, embedded with Tissue-Tek OCT and frozen on dry ice. Samples were then sectioned at 12-16μm using a Leica CM1850 cryostat.

### Lineage tracing

Lineage tracing was performed using *Msx1^CreERT2/+^; Rosa26R^LacZ/LacZ^* mice. Single cell lineage tracing was performed using *Msx1^CreERT2/+^; Rosa26R^Ai14/Ai14^* mice. Tamoxifen powder (T5648-1G, Sigma-Aldrich) was diluted in 10% ethanol, 90% corn oil (C8267, Sigma-Aldrich) at a concentration of 20mg/mL, vortexed and agitated overnight at 37°C. For harvest of embryonic time points, pregnant female mice were weighed prior to injection and tamoxifen was injected intraperitoneally at a concentration of 1mg/10g of the animal. High concentration of tamoxifen caused dystocia in pregnant female mice, hence for harvest of postnatal samples, tamoxifen powder was diluted in corn oil at a concentration of 75 ug/mL and the pregnant female was injected intraperitoneally with 150μL of the diluted solution. At least 3 mice across 2 litters were used per immunohistochemistry experiment. For single cell lineage tracing on embryonic hindbrain/cerebellum collected at E10.5, E14.5, E18.5 tamoxifen was administered at E9.5, while for embryonic telencephalon collected at E14.5, E18.5 tamoxifen was administered at E12.5.

### Chick embryo collection

Fertilized chicken eggs were obtained from Sun State Ranch (Sylmar, CA) and Petaluma Egg Farm (Petaluma, CA), and grown at 37°C for 18-20 hours to reach HH4-5.

### Cloning of expression vector

The pCI-FLAG-EZHIP construct was generated by PCR amplifying the human *EZHIP* sequence from the copGFP-EZHIP construct with an N-terminal FLAG tag. It was then cloned into the pCI-H2B-RFP vector using Asc1 and Pme1 restriction sites.

Primers:

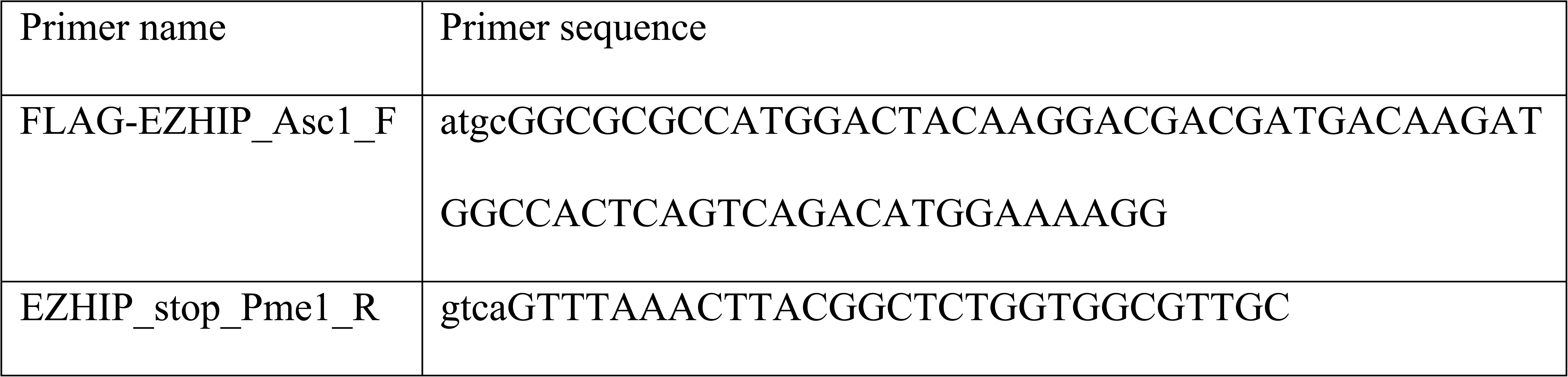

### Electroporation of chick embryos

*Ex ovo* electroporations were performed with stage HH4-5 embryos as described previously^2^. Embryos were dissected onto rings of filter paper in Ringer’s and a solution of DNA expression construct was injected into the space between the vitelline membrane and ectoderm. The DNA was then electroporated into the ectoderm with 5 pulses of 5.2V for 50ms, with 100ms between each pulse. Embryos were cultured at 37°C until HH9 or HH13, either in thin albumen with 1% penicillin/streptomycin alone or on a bed of agar albumin (50% thin albumin, 0.3% BactoAgar (214010, BD), 0.15% glucose, 61.5mM NaCl, 1% penicillin/streptomycin). All embryos were bilaterally electroporated with 2.5µg/µl of the control construct, pCI-H2B-RFP, on the left side and 2.5µg/µl of pCI-FLAG-EZHIP on the right side. Embryos were screened for RFP expression to ensure electroporation coverage and efficiency.

### Cell culture

Primary PFA cell lines were cultured as described previously^3^. Briefly, PFA cell lines were cultured in on laminin-coated (L2020, Sigma-Aldrich) Primaria plates (353803, Corning) pre-treated with poly-L-ornithine (A-004-M, Sigma Aldrich). Serum-free Human Neurocult NS-A Basal media (05750, StemCell Technologies) was supplemented with N2 (17502048, ThermoFischer Scientific), B27 minus vitamin A (12587010, ThermoFischer Scientific), EGF (E9644, Sigma-Aldrich), FGF (100-18B, Peprotech), heparin (H3393, Sigma-Aldrich), BSA (A8412, Sigma-Aldrich), Glutamine (35050061, Life Technologies) to make complete ependymoma media (EPN-C). All PFA cell lines were continuously cultured in hypoxic conditions (1% O2, 94% N2, 5% CO2). For 1% vs 21% O2 comparison PFA cell lines were cultured for 3 generations in 1% O2, 94% N2, 5% CO2 or 21% O2, 74% N2, 5% CO2, respectively.

### Flow cytometry and FACS sorting

PFA cell lines were treated with accutase (07922, STEMCELL), washed with 1xPBS and incubated with blocking buffer 1xPBS, 5% BSA for 10 min. Samples were then incubated with CXCR4-PE at a concentration of 1ug per 5 million cells (306506, BD Biosciences) in 1xPBS, 1%BSA for 20 min on ice in the dark. Samples were then washed with 1xPBS, 1%BSA 3 times and resuspended in 1xPBS, 1%BSA, 1:10000 DAPI solution and filtered through a 70μm filter. Samples were flow analyzed or flow sorted with MA900 and analyzed using FlowJo.

### *In vitro* Limiting Dilution Assay (LDA)

LDA was conducted in a 96 well Primaria plate (353072, Corning) coated with laminin as described above. PFA cell lines were stained with CXCR4-PE as described above and CXCR4+ and CXCR4- negative cells were sorted into a 96 well plate with EPN-C using single cell sorting of MA900. Exponentially increasing numbers of cells were seeded in wells with the highest number of 256 cells per well and 1 cell per well at the lowest dose with 4 technical replicates. Cells were fed every 4 days and scored for colonies after 1 month in culture. Statistical analysis was ran using Extreme Limiting Dilution Analysis (ELDA)^4^.

### Cryosectioning of chick embryos

Cryosectioning was performed to generate 18μm transverse sections of chick embryos using a Microm HM550 cryostat. Live or processed embryos were fixed in 4% paraformaldehyde overnight at 4°C, then incubated in 15% sucrose overnight at 4°C, followed by 7.5% gelatin overnight at 37°C prior to mounting in silicone molds and snap freezing in liquid nitrogen.

### Immunohistochemistry

#### Immunohistochemistry on mouse embryonic tissue

Sections were rehydrated with 1xPBS, blocked and permeabilized for 30 min with 5%BSA, 0.5% Triton-X, 1xPBS and incubated with primary antibodies for Msx1 (AF5045, R&D, 1:100), Cxcr4 (ab124824, Abcam, 1:500), Blbp (Ab32423, Abcam, 1:500), Gfap (Z0334, Agilent, 1:500), NeuN (Mab377, Millipore, 1:500), βgal (ab9361, Abcam, 1:2000), Sox10 (ab227680, Abcam, 1:100), Zfhx4 (HPA023837, Atlas, 1:200), Hoxb13 (90944, Cell Signaling Technology, 1:100), p73 (ab40658, Abcam, 1:500), H3K27me3 (9733S, Cell Signaling Technology, 1:500) for 2 hours at room temperature (Gfap and Sox10 antibodies were incubated overnight at 4°C) diluted in 1% BSA, 0.1% Triton-X, 1xPBS. After primary antibody incubation, sections were washed three times with 0.1% Triton-X, 1xPBS and incubated with secondary antibodies (Alexa Fluor 488, 594 or 647, Invitrogen) for 1 hour at room temperature. Samples were then washed three times and counterstained with DAPI (1:1000) for 10 minutes. Following DAPI incubation, samples were mounted with DAKO (Agilent, S3025). Samples were imaged using Leica SP8/STED confocal microscope. For Msx1/H3K27me3 staining, Z-stacks were made, and maximum intensity projection was created using Z-project in Fiji^5^.

#### Immunohistochemistry on mouse tumors

Formalin-fixed, paraffin-embedded sections were analyzed by Hematoxylin and Eosin or MSX1, GFAP and S100. In brief, tissue sections were incubated in Tris EDTA buffer (cell conditioning 1; CC1) at 95°C for 1 hour to retrieve antigenicity, followed by incubation with MSX1 (AF5045, R&D, 1:200), GFAP (ab68428, Abcam,1:250) or S100 (MA-32625, Invitrogen, 1:1000) antibody for 1 hour. Slides were then incubated with secondary antibody (Jackson Laboratories) with 1:500 dilution followed by Ultramap HRP and Chromomap DAB detection.

#### Immunohistochemistry on human embryonic tissue

Formalin-fixed-paraffin-embedded (FFPE) cerebellar sections were prepared at a thickness of 4 microns along the sagittal plane and mounted on Superfrost Plus slides from Thermo Fisher Scientific. To maintain antigenicity and prevent RNA degradation, the slides were kept refrigerated. IHC was carried out as previously described^6^. The primary antibodies used in the study included: MSX1 (AF5045, R&D Systems). Fluorescent dye-labeled secondary antibodies from ThermoFisher (Alexa Fluor 488 and 594, 1:1000) were employed. Following incubation with secondary antibodies, sections were counterstained with DAPI (4’,6-diamidino-2-phenylindole) using Vectashield mounting medium (Vector Laboratories, H-1200).

#### Immunohistochemistry on human tumors

Immunohistochemistry with an anti-MSX1 antibody (AF5045, R&D Systems) was performed on 5µm FFPE sections at a dilution of 1:1200. Antigen retrieval was performed in Dako Target Solution pH9 (S236784-2, Agilent) followed by an overnight primary antibody incubation. Sections were then treated with Goat-HRP Polymer (ab214881, Abcam), detected using NovaRed (SK-4800, Vector Laboratories), and counterstained with hematoxylin (H3404, Vector Laboratories). Formalin-fixed, paraffin-embedded sections were analyzed for PAX3 expression. In brief, tissue sections were incubated in Tris EDTA buffer (cell conditioning 1; CC1) at 95C for 1 hour to retrieve antigenicity, followed by incubation with PAX3 (LS-B3737 at 1:200) antibody for 1 hour. Slides were then incubated with secondary antibody (Jackson Laboratories) with 1:500 dilution followed by Ultramap HRP and Chromomap DAB detection. Intensity scoring was done on a common four-point scale. Descriptively, 0 represents no staining, 1 represents low but detectable degree of staining, 2 represents clearly positive staining, and 3 represents strong expression. Expression was quantified as H-Score, the product of staining intensity, and % of stained cells.

#### Immunohistochemistry on human autopsy samples

Histochemical staining of formalin-fixed paraffin embedded (FFPE) tissue samples was done with xylene de-paraffinization, ethanol rehydration and heat-induced epitope retrieval (HIER) carried out in a Tris-EDTA (TE) buffer of 9 pH in a pressure cooker. Permeabilization of slides was done using a phosphate buffered saline (PBS) and 0.25% Triton X-100 solution over a 15-minute incubation. Protein blocking was done using a PBS, 5% Bovine Serum Albumin (BSA) and 0.1% Tween-20 solution over a one-hour incubation. Overnight incubation at 4°C was done using anti-MSX1 antibody (AF5045, R&D Systems, 1:500) in a PBS, 5% BSA and 0.1% Tween-20 solution and slides were washed after incubation three times using a PBS and 0.1% Tween-20 solution. Peroxidase block consisting of 3% H2O2 for a 15-minute incubation was carried out and slides were washed again in PBS and 0.1% Tween-20 solution. A pre-diluted secondary horse anti-goat polymer (ImPRESS Horse-anti-goat HRP, Vector laboratories) was used for a 30-minute incubation and slides were washed again in PBS and 0.1% Tween-20. DAB solution (DAKO Omnis) was applied for approximately 2-3 minutes before washing and hematoxylin staining for one minute. Slides were then dehydrated in increasing concentrations of ethanol and mounted using coverslips. Imaging was performed using an Olympus light microscope.

#### Immunohistochemistry on chick embryos

HH13 embryos were fixed overnight at 4°C in 4% paraformaldehyde. After sectioning onto glass slides, they were blocked for 1 hour at room temperature in 10% goat or donkey serum in PBST. Primary antibody incubation occurred overnight for one night at 4°C in 10% goat or donkey serum, followed by three 10-minute washes at room temperature in PBST, then incubation in secondary antibody and 1:10,000 DAPI overnight at 4°C for one night in 10% goat or donkey serum. Slides were then washed in PBST three times for 10 minutes each and imaged. Primary antibodies used: Mouse IgG1 anti-Pax7 (Developmental Studies Hybridoma Bank, 1:10); Rb anti-FLAG (F7425, Sigma, 1:500); Ms anti-FLAG M2 (F1804, Sigma, 1:500); rabbit anti-Sox9 (AB5535, EMD Millipore, 1:1000); Rb anti-H3K27me3 (C36B11, Cell Signaling, 1:1000). Secondary antibodies used: Molecular Probes donkey or goat secondary antibody conjugated to Alexa Fluor 488, 568, or 647 (1:1000).

### *In situ* Hybridization Chain Reaction (HCR)

All HCR was performed with probes designed by Molecular Technologies and following the published V3 protocol^7^. A 9-probe set was used for chicken Msx1 and a 20-probe set was used for human EZHIP. Prior to sectioning, embryos were postfixed in 4% paraformaldehyde overnight at 4°C. After sectioning, slides were stained with 1:10,000 DAPI in PBS for 30 minutes at room temperature.

### RNAscope

#### Mouse embryos

Mouse cryosections were prepared according to ACD RNAscope Fixed Frozen Tissue technical note (320534) and stained according to ACD RNAscope 2.5 HD User Manual (322360-USM). Briefly, OCT sections were rehydrated with 1xPBS, post-fixed in cold 4% PFA, dehydrated in ethanol gradient. Following ethanol dehydration samples were incubated with RNAscope hydrogen peroxide solution for 10 min at room temperature. Afterwards, sections were incubated with Protease Plus for 10 minutes at room temperature. Samples were then incubated with the experimental probes Wnt3a (405041, ACD RNAscope), Wnt7b (401131, ACD RNAscope), followed by AMP probes and Fast Red staining. Multiplex fluorescence RNAscope was done according to ACD RNAscope Multiplex Fluorescent Reagent Kit v2 User Manual (323100-USM). Samples were incubated with the experimental probes Msx1 (421841, ACD RNAscope), Wnt1-C3 (401091-C3, ACD RNAscope) and Cxcr4-C4 (425901-C4, ACD RNAscope), followed by AMP probes, anti-C1, C3 and C4 HRP, and Opal Dyes 520 (1:1500, FP1487001KT, Akoya Biosciences), 620 (1:500, FP1497001KT, Akoya Biosciences) and 690 (1:500, FP1495001KT, Akoya Biosciences). Sections were then counterstained with Methyl Green (H-3402-500, Vector Laboratories) or DAPI (ACD RNAscope) and mounted with Ecomount (EM897L, Biocare Medical) or Dako (Agilent, S3025).

#### Human embryos

ISH assays were conducted using commercially available probes from Advanced Cell Diagnostics, following the manufacturer’s protocols without any modifications. The probes utilized in this study included: CXCR4 (310511, ACD RNAscope), EZHIP (1107811, ACD RNAscope). All sections were counterstained with hematoxylin or methyl green.

### 10x#single cell and single nucleus RNA sequencing

#### Preparation of samples for single cell lineage tracing

*Msx1^CreERT2/+^;R26R^Ai14/Ai14^*mice were injected with tamoxifen intraperitoneally at E9.5 for hindbrain/cerebellar harvesting and E12.5 for telencephalic harvest.

#### Preparation of single-cell suspensions

Fresh patient tumors were collected at the time of surgical resection. Tumor tissue was mechanically dissociated followed by collagenase-based enzymatic dissociation as previously described^8^. Embryonic samples were dissected as previously described and dissociated using Papain dissociation system (LK003150, Worthington Biochemical Corporation)^8^.

#### Single cell RNA library preparation and sequencing

Single cell suspensions were assessed with a trypan blue count. We aimed to load 10,000-15,000 cells per sample using the Chromium Controller in combination with the Chromium Single Cell 3’ V3 and V3.1 Gel Bead and Chip kits (10X Genomics). Individual cells were partitioned into gel beads-in-emulsion (GEMS), followed by reverse transcription of barcoded RNA and cDNA amplification. Individual single-cell libraries with indices and Illumina P5/P7 adapters were generated with the Chromium Single Cell 3’ Library kit and Chromium Multiplex kit. The libraries were sequenced on an Illumina Novaseq6000 sequencer.

#### Preparation of single-nuclei suspensions

Nuclei were isolated from fresh, snap frozen tumor tissues as previously described^9^. Frozen tissues were doused in 1 ml of chilled lysis buffer (lysis buffer; 10mM Tris-HCl (pH 7.4), 10mM NaCl, 3 mM MgCl2, 0.05% NP-40 detergent) 5 times with a loose pestle, 10 times with a tight pestle and lysed for 10 minutes on ice. 5 ml of chilled wash buffer (wash buffer; 5% BSA, 0.04U/µL Rnase inhibitor, 0.25% glycerol) was added to the sample, passed through a 40 µm cell strainer and centrifuged at 500 x g for 5 minutes at 4°C. After pelleting, the nuclei were resuspended in 5-10 ml of wash buffer. After two washes, single-nuclei suspensions were passed through a 20 µm cell strainer, pelleted, and resuspended in PBS with 0.05% BSA.

#### Single-nuclei RNA library preparation and sequencing

Single nuclei suspensions were assessed with a trypan blue count. We aimed to load 10,000-15,000 cells or nuclei per sample using the Chromium Controller in combination with the Chromium Single Cell 3’ V3 and V3.1 Gel Bead and Chip kits (10X Genomics). Individual cells or nuclei were partitioned into gel beads-in-emulsion (GEMS), followed by reverse transcription of barcoded RNA and cDNA amplification. Individual single-cell libraries with indices and Illumina P5/P7 adapters were generated with the Chromium Single Cell 3’ Library kit and Chromium Multiplex kit. The libraries were sequenced on an Illumina Novaseq6000 or Illumina Novaseq X Plus sequencer.

### Single cell and single nucleus RNAseq analysis

#### CellRanger

Filtered count matrices were generated using the 10X Genomics CellRanger (v5.0.1) pipeline. In brief, raw base call (BCL) files were demultiplexed with default settings and introns were included and resulting FASTQs were aligned to a reference genome. For single cell lineage tracing experiments reads were aligned to a custom reference of mouse genome with a tdTomato-WPRE-polyA transcript sequence, which was obtained from the previously published plasmid (Addgene plasmid #22799) used to create Ai9 and Ai14 (current study) mice.

#### Quality control, normalization and clustering

Quality control, normalization and clustering on all objects was performed using Seurat (v.4.3.0) according to the Seurat vignette and as previously published^8,10,11^. Briefly, quality of cells was assessed per sample, with low quality cells being excluded from the analysis based on mitochondrial gene content and gene counts. For single cell lineage tracing samples, tighter quality control was performed on nFeature_RNA counts. Density function was run on nFeature_RNA counts across all cells using bandwidth of 0.5. Local minima and maxima were calculated using turnpoints function in pastecs package (v_1.3.21). First pit value was used as the new minimum nFeature_RNA threshold since the first peak in the nFeature_RNA distribution would suggest lower quality of the cells. For maximum nFeature_RNA counts 5 times median absolute deviation of nFeature_RNA was used. The samples were merged, normalized using sctransform (v.0.3.5) and corrected for batch effects using harmony (v.0.1.1). To infer copy number variation in the tumor sc/snRNAseq objects, InferCNV (v.1.14.2) was used using ‘SCT’ assay of the sc/snRNAseq objects. Markers for each tumor cell cluster were determined using FindAllMarkers function in Seurat and cells expressing microglial or vasculature genes were used as a reference input for InferCNV. Malignant versus non-malignant cells were determined using a combination of InferCNV and marker expression. Cycling cells were removed from the PFA tumor subset.

#### Identifying tdTomato^+ve^ cells

Density function was run on tdTomato counts across all cells using bandwidth of 0.5. Local minima and maxima were calculated using turnpoints function in pastecs package (v_1.3.21). The first peak of expression was recognized as background signal of tdTomato. First and fourth pit values were used as the new minimum tdTomato threshold for hindbrain and telencephalic samples, respectively, with cells expressing tdTomato levels higher than this threshold called ‘tdTomato+ve’. For subsetting Msx1-derived lineages, clusters with more than 10% (for hindbrain) and 5% (for telencephalon) tdTomato+ve cells were subsetted out of the integrated objects per timepoint and reclustered. Initial annotations were preserved in the subsetted dataset.

#### Differentiation potential and pseudotime analysis

To perform differentiation potential analysis, we ran the CytoTRACE package (v.0.3.3) on SCT@counts slot of PFA-EPN object^12^. To reconstruct the pseudotime trajectory we utilized the slingshot package (v.2.6.0) using NSC cluster as a ‘root’^13^.

#### Differential gene expression heatmap

Differentially expressed genes across clusters for the PFA, mouse and human embryonic datasets were identified using FindAllMarkers function in Seurat. Genes were then ordered based on avg_log_2_FC. Single cell matrix was then subsetted based on which genes were found in the scale.data slot in the SCT assay. The matrix was then plotted using Heatmap function from ComplexHeatmap package (v.2.14.0).

#### Analysis of regulon activity

The activity of specific regulons in each tumor cell type was inferred using the package pySCENIC (v.0.10.0), implemented in python (v.3.7.6), with the parameter - min_genes 10. For each cluster, SCENIC AUCell score matrix was scaled and top 10 regulons per tumor cell type were plotted using ComplexHeatmap package (v.2.14.0).

#### Pre-NC/RP annotation

Top 70 genes sorted by avg_log2FC were used from Pre-migratory NCCs I and II annotated cells from Dong *et al.* dataset and cells within PFA ependymoma scRNAseq object were scored using AddModuleScore function of Seurat (v.4.3.0). Pre-NC/RPs were then annotated using WhichCells function. To annotate the cells, the score had to meet the following conditions:

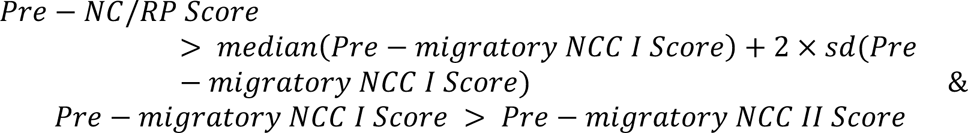

#### Transcriptional mapping

To compare PFA ependymoma to both mouse and human embryonic datasets, we utilized CHETAH package (v.1.14.0). For the mouse dataset gorth function from gprofiler2 package (v.0.2.3) was used to convert *M.musculus* genes to *H.sapiens*. To construct input and embryonic references RNA@data slot was utilized. In addition, to construct references from embryonic datasets, we sampled 200 cells per cluster 5 times and ran CHETAHclassifier function with threshold of 0 to force mapping of all cells. Average percentage mappings from 5 CHETAH iterations per PFA cell type were amalgamated to generate annotations. For single cell lineage tracing cluster scores in PFA cell types, top 100 differentially expressed genes from each embryonic cluster was compiled into a respective module score using AddModuleScore function in Seurat. Scores were then plotted across PFA cell types using DotPlot function in Seurat.

### Analysis of previously published datasets

#### Single Cell RNA sequencing

Data from Dong *et al*., 2020 and Gojo *et al*., 2020 was downloaded from GSE137804 and GSE141460 respectively^15,16^. Only cells related to neural tube and neural crest were subsetted from the Dong *et al*., including Pre-migratory NCCs I, Pre-migratory NCCs II, Neural Tube Cells, Neural Crest Cells, Migratory Neural Crest Cells, Melanocytes and Sensory Neurons. Based on previously published markers, Pre-migratory NCCs I and II were renamed as Pre-NC/RPs and Roof Plate Stem Cells, respectively. Normalization and clustering on the Dong *et al*. data was performed according to Seurat vignette. The samples were merged, normalized using sctransform (v.0.3.5) and corrected for batch effects using harmony (v.0.1.1). Gojo *et al*. dataset was normalized and clustered according to Seurat vignette. The samples were merged, normalized using sctransform (v.0.3.5). Gojo *et al*. dataset was not corrected for batch effects. To plot the Pre-NC/RPs signature in Gojo *et al*. dataset, top 70 genes sorted by log2FC were used from Pre-NC/RPs cluster in Dong *et al*. as described previously. The DotPlot with the Pre-NC/RPs signature expression was created using dittoDotPlot function from the dittoSeq package (v.1.10.0).

#### Bulk RNA sequencing

Previously published bulk RNA sequencing of PFA-EPN and ZFTA-EPN from Michealraj *et al*., 2020 (EGAD00001006046) was reanalyzed with respect to unique markers of ZFTA-EPN^3^. Volcano plot was generated using geom_point function from ggplot2 package (v.3.4.2).

#### Microarray gene expression profiling

Microarray data from Pajtler *et al*., 2015 (GSE64415) was accessed used R2 genome browser^17^. Expression data for the probes of interest was downloaded and plotted using Prism. Expression of *ZFHX4* was analyzed using Mann-Whitney test. Expression of *KIT* and *GRIN3A* was analyzed using Kruskal-Wallis test.

#### Midline vs. lateral PFB-EPN

Patient data was downloaded from Fukuoka *et al.* (2018)^18^. Tumors annotated as PFA-EPN or PFB-EPN were segregated by annotated primary tumor location, and stacked bar plot was made using ggplot2 package (v3.4.2).

#### Anatomic distribution of ZFTA-EPN

Patient data was downloaded from Pohl *et al*., 2024^18^. Tumours with predicted subgroup of ZFTA-EPN and a known location were selected and counted. Pie chart was made using ggplot2 package (v.3.4.2).

### Chick embryo imaging analysis

All images were obtained using either a Zeiss AxioImager.M2 with an Apotome.2 or a Nikon CSU-W1. Cell counts of PAX7 and Sox9 positive cells in sections of the cranial region were performed using the Analyze Particles feature in Fiji^5^. PAX7 positive cells remaining in the neural tube and Sox9 positive cells present in the pharyngeal arches were counted in HH13 embryos. For PAX7 cell counts at HH13, 7-8 non-adjacent images spanning pharyngeal arch level to first somite (∼every sixth section) were quantified and averaged per embryo. Experimental values were divided by control values from the same image. For Sox9 cell counts in the pharyngeal arches at HH13, we quantified 5-10 non-adjacent images (taking every 2nd or 4th section depending on embryo size and number of sections throughout). Control and experimental cell counts in the arches were summed per embryo to account for variation in the angle of each section through the pharyngeal arches, and total experimental cell count was divided by total control cell count for each embryo. Fluorescence intensity of Msx1 HCR signal and H3K27me3 immunofluorescence signal was measured from HH13 sections by manually defining regions of interest to measure integrated density, subtracting the per pixel background signal averaged from three manually defined background regions, then normalizing to DAPI signal from the defined regions of interest. The regions of interest included experimental and control halves of the neural tube, from dorsal midline to ventral midline for both MSX1 and H3K27me3, as well as control and experimental non-neural ectoderm and otic placode for H3K27me3. 5-15 non-adjacent images spanning pharyngeal arch level to first somite (roughly every 3rd-4th section) were quantified and averaged per embryo for all regions except otic placode, for which only 1-2 sections were analyzed per embryo due to the size of the placode. Experimental values were divided by control values from the same image. For all statistical analysis on images, a paired two-tailed Student t test was performed to compare two values (experimental and control) within single embryos.

### Data Availability

Mouse hindbrain scRNAseq data has been deposited to GSE262208. Raw sequencing data for PFA-EPN has been deposited to EGA50000000478 and CNP0005408. Previously published PFA-EPN scRNAseq has been deposited to EGAS00001003170. Raw sequencing data from single nucleus RNAseq of PFB-EPN, MPE-EPN and scRNAseq of ZFTA-EPN and mouse lineage tracing are being uploaded to EGA. The accession numbers are pending.

